# Prime editing of an evolutionarily informed PEPC residue enhances photosynthesis and grain nutrition in C_3_ rice

**DOI:** 10.64898/2026.01.14.699509

**Authors:** Sonali Panda, Tanmoy Halder, Chandana Ghosh, Manaswini Dash, Debasmita Panda, Subhasis Karmakar, S. P. Avinash, Deeptirekha Behera, Tushar Mandol, Totan Adak, Rubina Khanam, Arabinda Mahanty, Anusha Thaikkandiyil, Sagar Banerjee, Krishnendu Chattopadhyay, Ajay Kumar Pandey, Swapan K. Datta, Kalavathi M. Kumar, Mridul Chakraborti, Meera K. Kar, Mirza J. Baig, Kutubuddin A. Molla

**Affiliations:** ICAR-Central Rice Research Institute, Cuttack-753006, India; Ravenshaw University, Cuttack-753003, India; ICAR-National Institute of Natural Fibre Engineering and Technology, Kolkata-700040, India; Department of Bioinformatics, School of Life Science, Pondicherry University, Kalapet-605014, India; National Agri-Food Biotechnology Institute, Sector 81, Knowledge City, S.A.S. Nagar, Punjab-140306, India; School of Agriculture, Swami Vivekananda University, Barrackpore, West Bengal-700121, India

## Abstract

C_4_ photosynthesis evolved through adaptive changes in key metabolic enzymes that enhance carbon fixation efficiency, yet transferring such traits into C_3_ crops has met limited success. Using precision prime editing, we introduced an evolutionarily conserved single amino acid substitution (Arg→Gly) into phosphoenolpyruvate carboxylase (PEPC) in rice. Edited plants exhibited enhanced PEPC activity in the presence of the feedback inhibitor malate, increased chlorophyll content, elevated photosynthetic rates under ambient CO_2_, and increased seed size and weight. Remarkably, seeds from the edited lines showed simultaneous enrichment of zinc (Zn), iron (Fe), and protein, hereafter referred to as ZiP rice. This work demonstrates that a single, evolutionarily guided residue change can reprogram core carbon metabolism in a C_3_ crop, enabling concurrent improvements in photosynthesis and grain nutritional quality.

## Main Text

Enhancing photosynthetic efficiency in C_3_ crops is a key objective in efforts to meet the increasing global food demand. One promising avenue is the introduction of C_4_-like traits into C_3_ crops, particularly rice, to boost carbon assimilation, yield potential, and climate resilience. Efforts to engineer C_4_ traits into C_3_ rice have met with limited success (*1*–*4*). Phosphoenolpyruvate carboxylase (PEPC), a key enzyme of the C_4_ pathway, catalyses the carboxylation of phosphoenolpyruvate to oxaloacetate and is tightly regulated by metabolites such as malate. Acquisition of greater PEP substrate saturation constants and increased tolerance towards feedback inhibition are crucial achievements in the evolution of C_4_ PEPC from the C_3_ ancestor (*5*). Comparative analyses reveal that a single amino acid change (R883G) in C_4_ PEPC reduces malate-mediated feedback inhibition (*6*). We wondered whether altering this residue in the endogenous rice PEPC to enhance its activity would impact metabolite balance—particularly PEP, oxaloacetate, malate, and citrate—thereby influencing rice physiology.

Prime editing, a precision genome editing technology, enables the introduction of targeted single-nucleotide changes without inducing double-strand breaks or permanently integrating foreign DNA (*7, 8*). Unlike traditional transgenic approaches, which face regulatory and public acceptance challenges (*9*), prime editing allows installation of evolutionarily informed variations that confer C_4_-like properties to C_3_ PEPC.

In this study, we employed prime editing to introduce a single evolutionarily guided amino acid substitution (Arg→Gly) in C_3_ PEPC in rice as a model. The edited lines exhibited enhanced PEPC efficiency in the presence of malate, increased photosynthetic rate, and enlarged seed size. Notably, seeds from edited lines were biofortified with significantly higher zinc, iron, and protein (ZiP) content, providing an attractive strategy to combat malnutrition. Our findings demonstrate that a single, evolutionarily informed residue change in PEPC can simultaneously enhance photosynthesis, agronomic performance, and nutritional quality in C_3_ crops, offering a powerful approach for multidimensional crop improvement.

### R883G amino acid substitution in rice endogenous PEPC reduces malate binding affinity

Comparative sequence analysis between C_3_ and C_4_ PEPC isoforms revealed several amino acid substitutions associated with the evolutionary transition from C_3_ to C_4_ photosynthesis. Notably, a substitution at position 883 was identified as a key determinant of the marked difference in inhibitor sensitivity between the two types (*6*). Multiple sequence alignment showed that PEPC isoforms from C_4_ plants carry a glycine (G) residue at this position, whereas the corresponding C_3_ isoforms retain an arginine (R) (Fig. 1A-B and table S1). The residue Arg883 resides within the feedback inhibitor-binding pocket of C_3_ PEPC (*6*). To evaluate whether the R883G substitution in rice PEPC alters inhibitor-binding chemistry, we performed molecular docking analysis using OsPEPC_R883_ and OsPEPC_G883_ with malate as the inhibitor. Docking results confirmed the involvement of the R883 residue in malate binding by OsPEPC. Although R883 does not directly interact with the inhibitor, it functions as a molecular clamp by forming an additional hydrogen bond that prevents the dissociation of malate from the binding site. In contrast, Gly883 in OsPEPC_G883_ is positioned farther from the inhibitor molecule, providing greater steric freedom that facilitates its release from the binding pocket (Fig. 1C-1H and fig. S1 A-B). These findings indicate that substitution of arginine with glycine at position 883 in rice PEPC could reduce malate sensitivity.

**Fig. 1.**
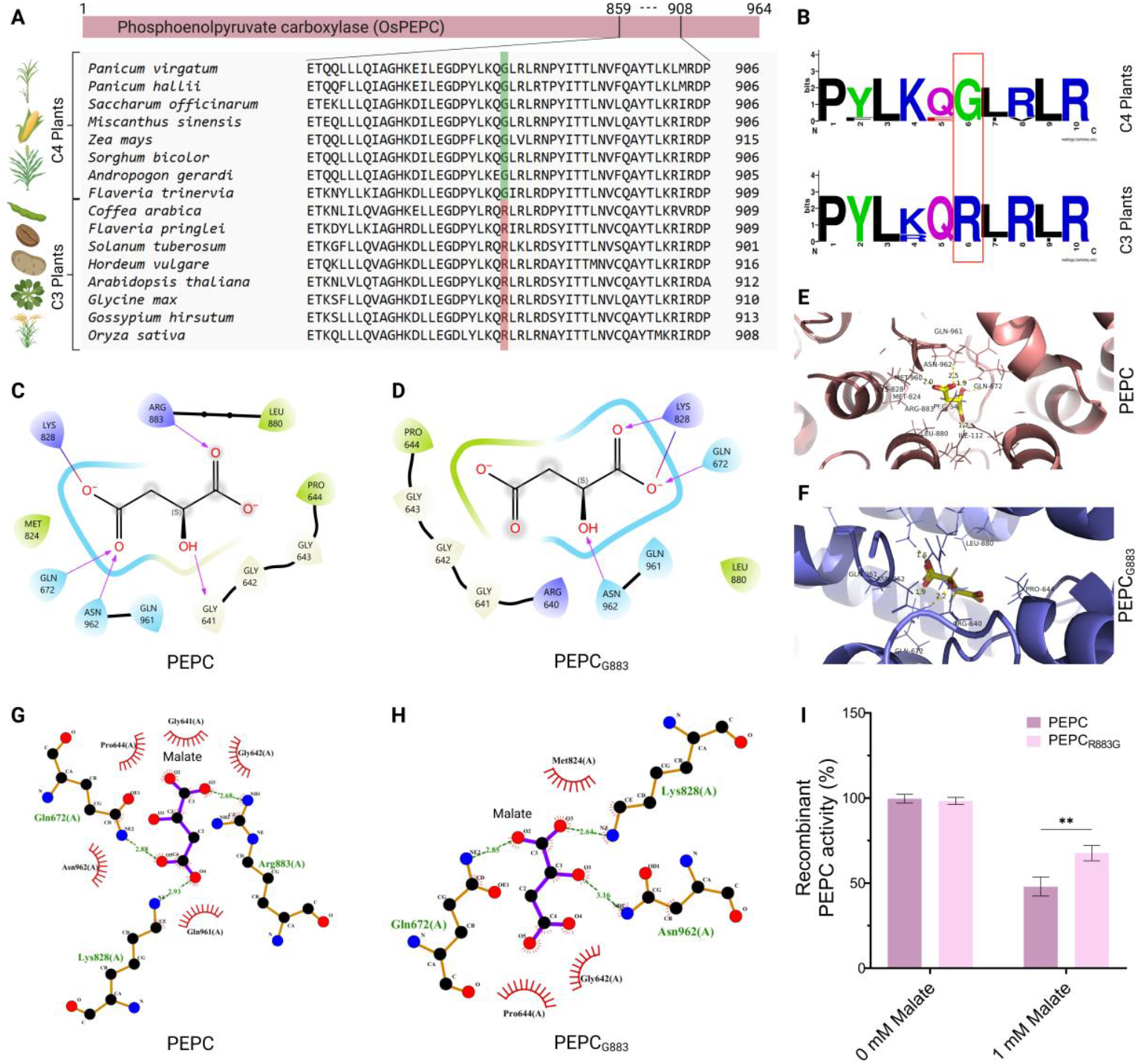
Evolutionary conservation, structural basis, and functional impact of the Arg→Gly substitution in OsPEPC. (**A**) Multiple sequence alignment of the C-terminal region of phosphoenolpyruvate carboxylase (PEPC) from representative C_4_, and C_3_ plant species. Evolutionarily conserved residues are highlighted in green for C_4_ and red for C_3_ species. (**B**) Sequence logo representations showing the conservation of the targeted motif in C_4_ plants (upper panel) and C_3_ plants (lower panel). Red box highlights the altered amino acid during evolution from C_3_ to C_4_. (**C, D**) Two-dimensional interaction maps depicting malate binding within the PEPC active site of the wild-type OsPEPC and mutated OsPEPC_R883G_ proteins. R883G substitution changes the hydrogen bonding pattern and electrostatic interactions. (**E, F**) Three-dimensional structural models of OsPEPC showing the malate-binding pocket and surrounding residues in the wild-type and mutated proteins. (**G, H**) Detailed ligand–protein interaction diagrams highlighting altered contacts between malate and OsPEPC residues in the wild-type and mutated (OsPEPC_R883G_) forms. Arg883 supports a tight binding of malate in wild-type OsPEPC. (**I**) Recombinant PEPC activity (%) of wild-type and mutated OsPEPC_R883G_ proteins measured in the absence (0 mM) and presence (1 mM) of malate. Data are presented as mean ± SD, and statistical significance is indicated by **P < 0.01.

### Recombinant PEPC with R883G substitution is less sensitive to malate

To test whether the R883G substitution reduces malate sensitivity, we constructed expression vectors encoding the wild-type and mutant (PEPC_R883G_) coding sequences, and expressed them heterologously in *E. coli* for functional characterization. The wild-type PEPC coding sequence was amplified from rice leaf cDNA, and the PEPC_R883G_ variant was generated via site-directed mutagenesis and cloned into a bacterial expression vector. Upon induction, both constructs produced a ∼120 kDa protein band (PEPC-6×His) predominantly in the soluble fraction, as observed in SDS–PAGE (fig. S2A). We then purified the proteins using Ni–NTA affinity chromatography (fig. S2B) and confirmed their identity by immunoblotting with an anti-His antibody (fig. S2C).

Malate inhibition assays revealed that PEPC_R883G_ retained 38% higher enzymatic activity in the presence of 1 mM malate compared to the wild-type enzyme (Fig.1I). These results demonstrate that the R883G substitution confers reduced malate sensitivity, thereby enhancing PEPC catalytic performance under feedback-inhibited conditions.

### Standardized prime editing enables installation of R883G in endogenous rice PEPC

Encouraged by the results from recombinant PEPC analysis, we sought to introduce the R883G mutation into the endogenous rice PEPC gene via prime editing. Initially, we employed the first-generation prime editing vector, pK-PE2, with canonical prime editing guide RNA (pegRNA) and regenerated over 250 plants, but none carried the desired edit.

We then optimised the editing reagents using our established rice protoplast system (*10*). Three plant prime editing vectors—pK-PE2, pK-ePE2, and enpPE2—were tested, each carrying either pegRNA or engineered pegRNA (epegRNA) designed for the target PEPC mutation (C2647→G2647, resulting in Arg883→Gly883) (Fig. 2A-C). The epegRNA contains an evopreQ1 RNA motif, which stabilizes the 3′ end and enhances editing efficiency (*11*). Transfection efficiency was monitored using a Ubi: GFP plasmid, showing ∼85% efficiency at 18 h post-transfection (Fig. 2D). At 72 h post-transfection, genomic DNA was harvested for targeted PCR amplification and next generation sequencing (NGS) analysis. Between 170,000 and 250,000 reads were obtained per sample. Editing efficiencies varied among the three vectors, with enpPE2_epegRNA achieving the highest efficiency (6.5%) for the desired C-to-G substitution (Fig. 2E and fig. S3).

**Fig. 2.**
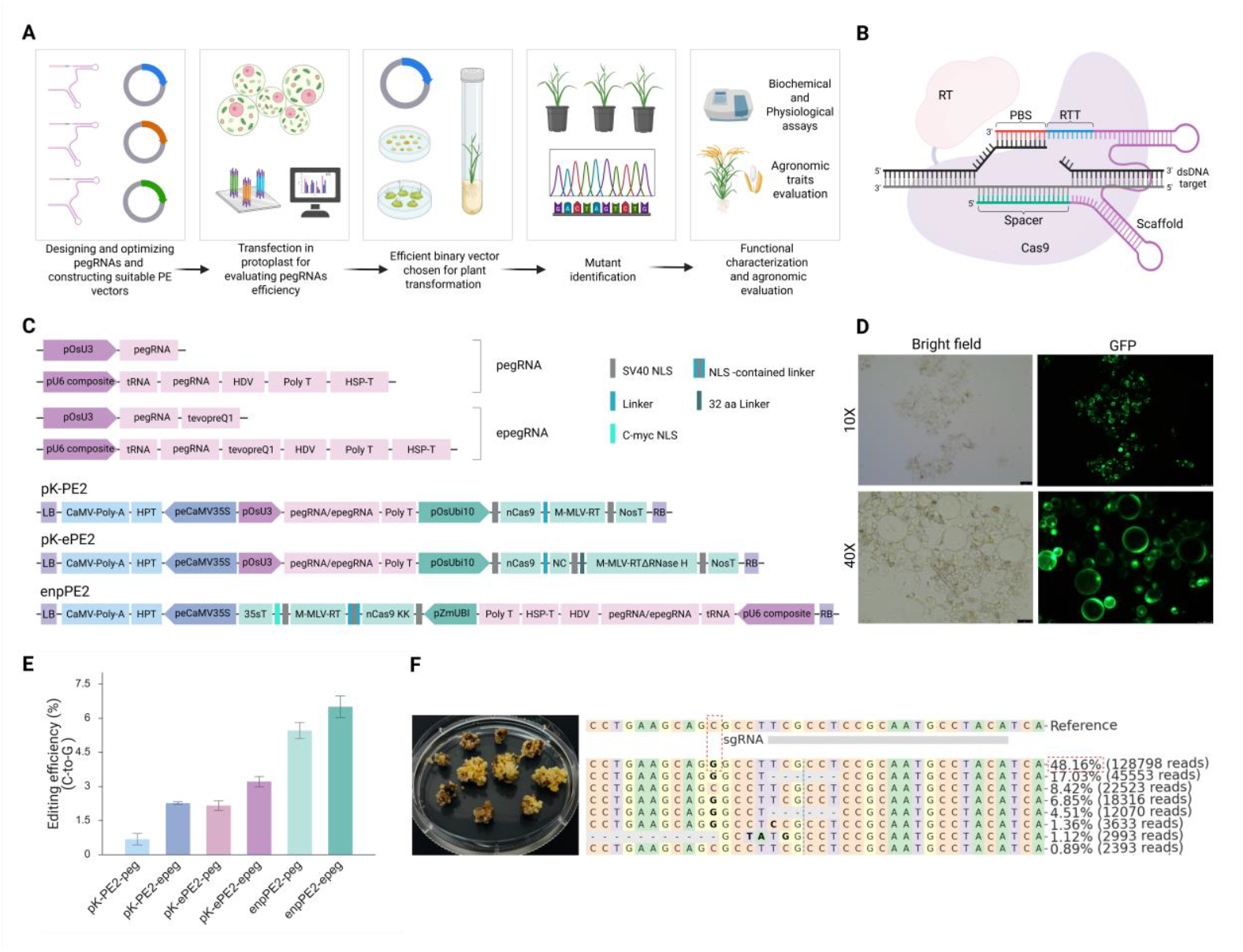
Establishment and optimization of prime-editing systems for precise R883G installation in OsPEPC. (**A**) Schematic overview of the prime-editing pipeline. (**B**) Schematic illustrating the prime editor–pegRNA complex bound to the target DNA locus. (**C**) Architecture of pegRNA and epegRNA expression cassettes and schematic maps of prime-editing vectors (pK-PE2, pK-ePE2, and enpPE2), highlighting promoters, terminators, selectable markers, Cas9 nickase, and RNA components used for plant transformation. Abbreviations: pOsU3/pOsU6, rice U3/U6 small nuclear RNA promoters; HSP-T, heat-shock protein terminator; Poly T, RNA polymerase III terminator; HDV, Hepatitis Delta Virus ribozyme; RT, reverse transcriptase; M-MLV-RT, Moloney Murine Leukemia Virus reverse transcriptase; nCas9, Cas9 nickase (H840A); NLS, nuclear localization signal; SV40 NLS, Simian Virus 40 nuclear localization signal; c-Myc NLS, c-Myc–derived nuclear localization signal; CaMV35S, Cauliflower Mosaic Virus 35S promoter; HPT, hygromycin phosphotransferase selectable marker; NOS-T, nopaline synthase terminator; LB/RB, left and right T-DNA borders. (**D**) Representative bright-field and GFP fluorescence images of rice protoplasts used to assess transfection efficiency. (**E**) Editing efficiencies (%) obtained using different prime-editing constructs, demonstrating enhanced performance of engineered enpPE2-epegRNA systems compared with other vectors (Data are presented as mean ± SD). (**F**) Selected calli carrying edited alleles and representative sequencing reads confirming precise nucleotide substitutions at the target site, with frequencies of edited reads indicated.

Encouraged by the protoplasts transient assay results, we performed *Agrobacterium*-mediated transformation using enpPE2_epegRNA to generate stable rice plants. Hygromycin-resistant calli were selected and screened via targeted deep sequencing. Consistent with protoplast data, enpPE2_epegRNA achieved ∼48% editing efficiency in selected rice calli (Fig. 2F), demonstrating robust activity in both transient and stable systems.

### R883G substitution boosts PEPC activity without altering transcription

We have regenerated 244 putative transgenic plants from hygromycin-selected calli. 93 lines were initially screened for the desired C-to-G edit by Sanger sequencing, revealing that 67 carried the targeted substitution—achieving 72% editing efficiency (Fig. 3A-B). Among these, 4 lines were biallelic and 63 monoallelic. Some plants also displayed pegRNA scaffold-derived byproducts (fig. S4). To obtain transgene-free homozygous lines, T_0_ plants were self-pollinated to produce T_1_ progeny. From the T_1_ generation, 45 plants were identified as homozygous for the desired edit, at which point screening was halted. PCR analysis targeting the RT and Cas9 sequences confirmed that 21 of these lines were transgene-free (fig. S5), from which five independent lines (3-4, 4-1, 23-7, 50-5, and 63-4) were randomly selected for further analysis.

**Fig. 3.**
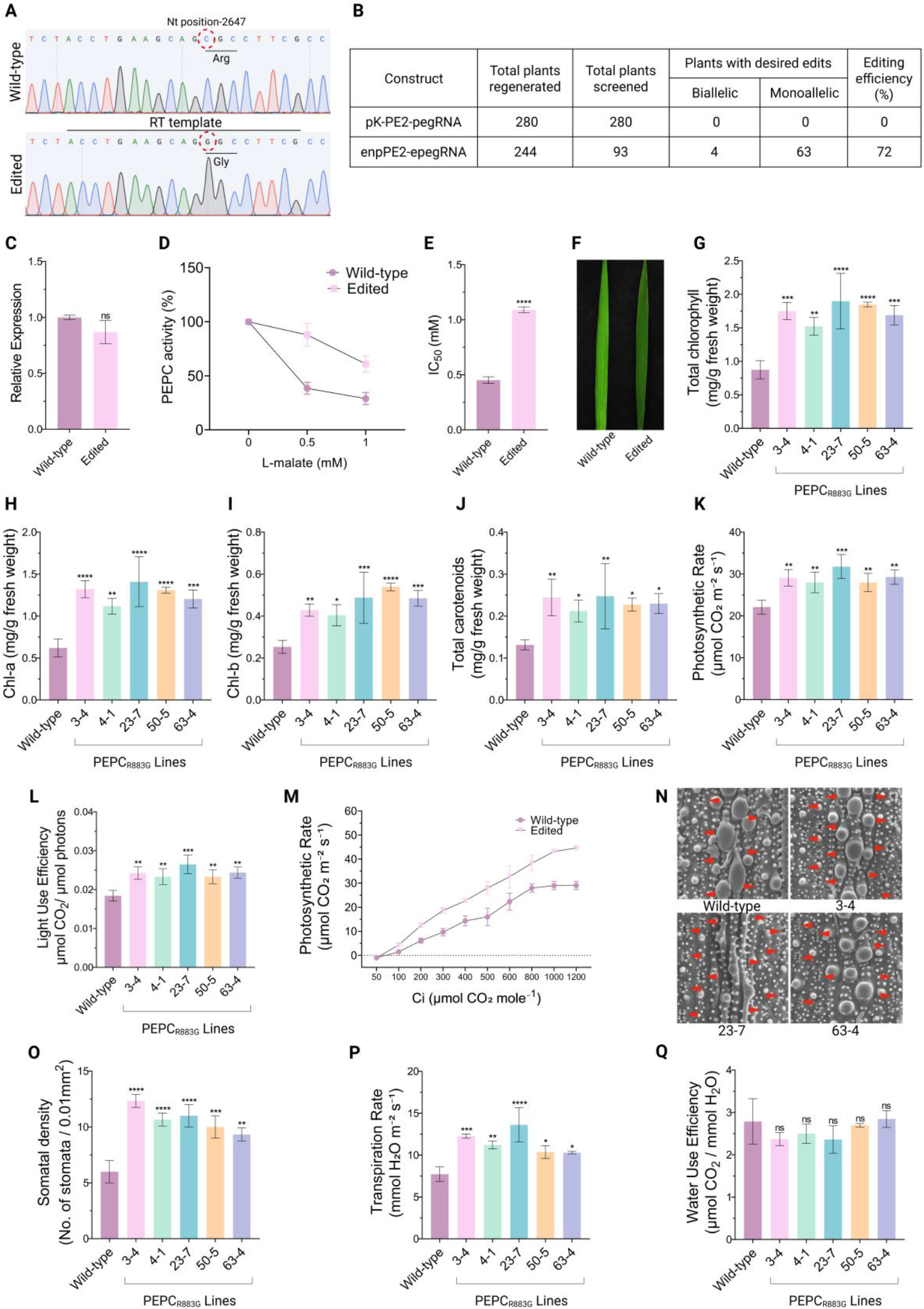
Impact of Prime editing–mediated modification of PEPC in rice physiology and biochemistry. (**A**) Representative Sanger sequencing chromatograms showing the targeted nucleotide substitution in *OsPEPC* at position 2647; the wild-type codon encodes arginine (Arg), whereas the edited lines carry a glycine (Gly) substitution, as indicated by red dashed circles. (**B**) Summary table showing regeneration and screening outcomes for different prime-editing constructs. (**C**) Relative expression of PEPC in wild-type and edited plants determined by qRT–PCR (ns: not significant). (**D**) PEPC enzyme activity (%) measured in the presence of increasing concentrations of L-malate. (**E**) IC_50_ values for L-malate inhibition of PEPC activity in wild-type and edited plants. (**F**) Representative leaves from wild-type and edited plants showing differences in greenness. (**G**) Total chlorophyll content (mg g^−1^ fresh weight). (**H**) Chlorophyll a (Chl a) content (mg g^−1^ fresh weight). (**I**) Chlorophyll b (Chl b) content (mg g^−1^ fresh weight). (**J**) Total carotenoid content (mg g^−1^ fresh weight). (**K**) Net photosynthetic rate (μmol CO_2_ m^−2^ s^−1^). (**L**) Light-use efficiency (μmol CO_2_ μmol^−1^ photons). (**M**) Photosynthetic rate at different intercellular CO_2_ concentration (Ci) at constant light intensity (1200 μmol m^−2^ s^−1^). (**N**) Scanning electron microscopy images of leaf epidermis showing stomatal distribution (red arrows) in wild-type and representative edited lines (0.01 mm^2^ area). (**O**) Stomatal density (number of stomata per 0.01 mm^2^). (**P**) Transpiration rate (mmol H_2_O m^−2^ s^−1^). (**Q**) Water use efficiency (μmol CO_2_ /mmol H_2_O). Data are presented as mean ± SD, and statistical significance relative to wild-type is indicated by asterisks (*P < 0.05, **P < 0.01, ***P < 0.001, ****P < 0.0001).

Potential off-target sites with high homology to the epegRNA spacer were examined in these five transgene-free lines. No off-target alterations were detected, confirming the precision and specificity of the editing system under our experimental conditions (fig. S6 A-C and table S3).

With editing accuracy and homozygosity confirmed, we assessed PEPC mRNA expression by qRT-PCR. Expression levels in edited lines (PEPC_R883G_) were comparable to wild-type (WT) (P = 0.2221; Fig. 3C), indicating that the mutation does not affect transcription.

We then evaluated the functional impact of the R883G substitution *via* malate inhibition assay. In the presence of 0.5 mM and 1 mM malate, edited lines exhibited 49.4% and 32% reductions in malate sensitivity, respectively (Fig. 3D), consistent with recombinant PEPC analysis. The IC_50_ of the modified PEPC was 1.09 mM malate, compared to 0.452 mM in wild-type, reflecting a ∼2.4-fold increase in malate tolerance (Fig. 3E). These findings align with observations in C3 *Flaveria pringlei*, where the analogous Arg-to-Gly substitution *in vitro* conferred a ∼3-fold enhancement in inhibitor tolerance (*6*). Collectively, these results demonstrate that a single, evolutionarily guided amino acid substitution markedly improves PEPC tolerance to malate inhibition without affecting transcription.

### PEPC_R883G_ rice plants accumulate more chlorophyll and exhibit enhanced photosynthetic capacity

Plants carrying the R883G substitution showed a substantial increase in photosynthetic pigment accumulation relative to wild-type. Total chlorophyll content increased by ∼2-fold, driven by ∼2-fold higher chlorophyll a and ∼1.8-fold higher chlorophyll b (Fig. 3F-I). Carotenoid levels were similarly elevated, showing an average 1.76-fold increase (Fig. 3J). A comparable enhancement in chlorophyll accumulation has been reported in rice lines overexpressing C_4_-type PEPC (*3*).

Gas-exchange measurements using an IRGA system revealed a clear enhancement in photosynthetic performance in the edited lines. The net CO_2_ assimilation rate (A) increased by ∼32% relative to wild-type, accompanied by a ∼32% improvement in light-use efficiency (Fig. 3K-L). A/Ci response analyses further demonstrated higher carboxylation efficiency and a consistently improved photosynthetic response across intercellular CO_2_ gradients, underscoring a robust enhancement in carbon-fixation capacity in PEPC_R883G_ plants (Fig. 3M).

Strikingly, PEPC_R883G_ plants also exhibited a marked increase in stomatal density (average 6 in WT vs. ∼10 in edited lines per unit area) (Fig. 3N-O), along with higher stomatal conductance (+53.4%) and transpiration rate (+49.6%) compared with wild-type (fig. S7A and Fig. 3P). Elevated PEPC activity is known to modulate stomatal aperture and transpiration through altered malate and related metabolite fluxes (*12*). Increased stomatal density and conductance, in turn, can enhance CO_2_ diffusion into the leaf, thereby reinforcing the observed gains in photosynthetic carbon assimilation (*13*).

Despite these changes, both water-use efficiency (WUE) and instantaneous WUE (iWUE) remained comparable between edited and wild-type plants, consistent with the proportional increases in CO_2_ assimilation and transpiration (Fig. 3Q and fig. S7B). These results suggest that improved PEPC activity—achieved here through a single amino acid substitution—can enhance photosynthetic pigment biosynthesis and strengthen overall photosynthetic performance.

### Prime-edited PEPC enhances iron and zinc accumulation in rice grains

Because PEPC_R883G_ plants displayed a dark-green phenotype with elevated chlorophyll content (Fig. 3F-I) and given that iron plays a central role in chlorophyll biosynthesis and chloroplast ultrastructure (*14*), we hypothesized that iron accumulation might also be altered in the edited plants. Consistent with this idea, Perls’ Prussian blue staining revealed a noticeably stronger blue coloration in seeds from edited plants compared with WT (Fig. 4A). Next, we quantified iron levels in unpolished and polished (1 min) grains using atomic absorption spectroscopy (AAS). Total iron content increased by 22.4% in unpolished grains (26.63 mg kg^−1^ in edited lines versus 21.75 mg kg^−1^ in WT), while polished grains from edited plants exhibited a 73.7% increase (20.24 mg kg^−1^ in edited versus 11.65 mg kg^−1^ in WT) relative to wild-type (Fig. 4B). This pronounced enrichment in polished grains indicates that iron accumulation is substantially enhanced within the endosperm of the edited plants. Notably, iron accumulation in polished grains of the prime-edited lines was comparable to or exceeded that reported for transgenic ferritin rice (∼18.89 mg kg^−1^) developed in the same Kitaake background (*15*).

**Fig. 4:**
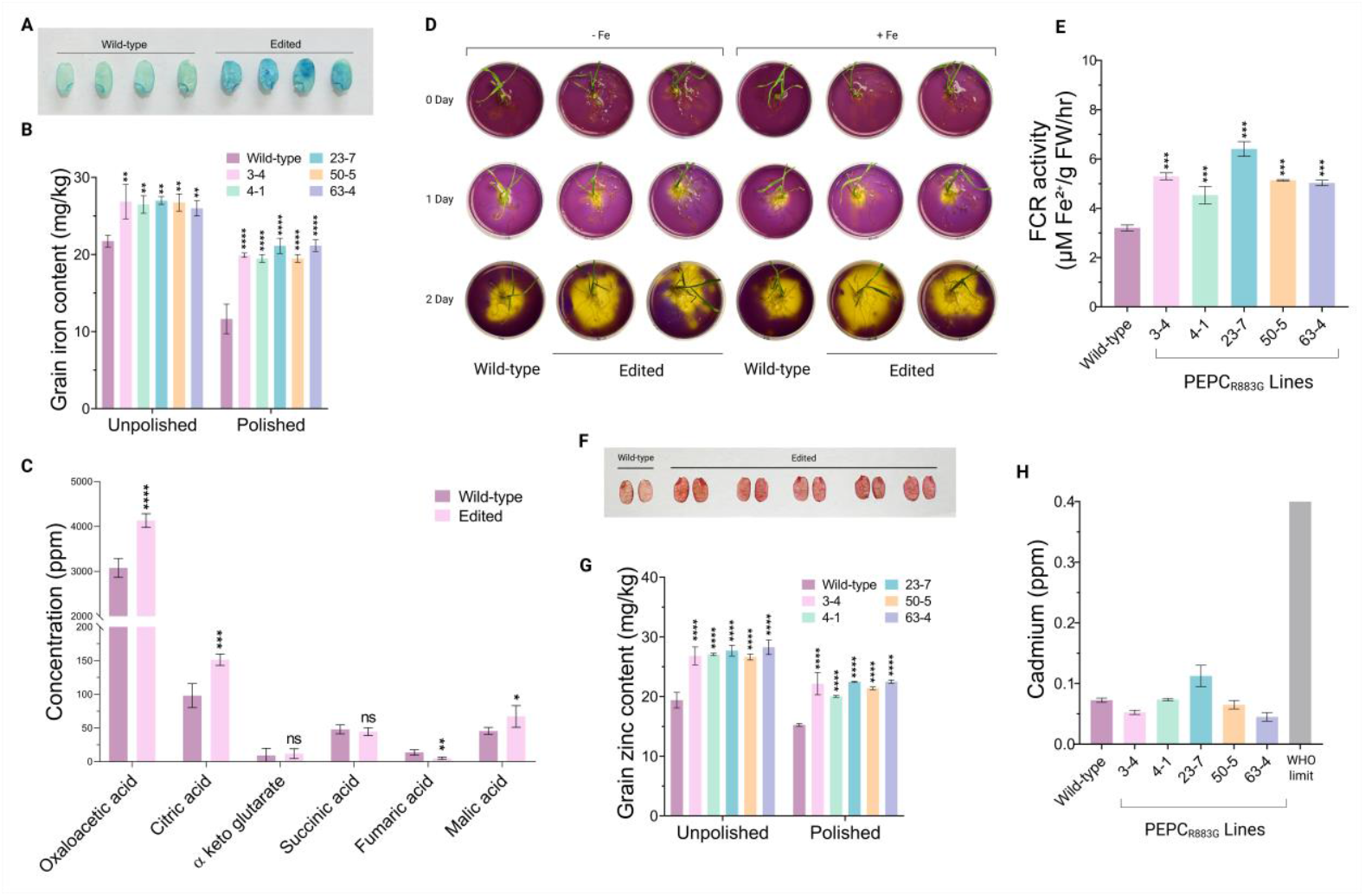
Enriched micronutrients in PEPC_R883G_ rice lines. (**A**) Representative image of Prussian blue–stained grains showing iron localization. (**B**) Grain iron content (mg kg^−1^) in unpolished and polished rice from wild-type and edited lines (Line-3, Line-4, Line-23, Line-50, Line-63). (**C**) Concentrations of TCA cycle intermediates in wild-type and edited plants. (**D**) Rhizospheric acidification assay comparing wild-type and edited rice seedlings under iron-deficient (–Fe) and iron-sufficient (+Fe) conditions. Yellow coloration indicates rhizosphere acidification resulting from proton extrusion by roots. (**E**) Ferric chelate reductase (FCR) activity in roots of wild-type and edited lines. Roots (100 mg fresh weight) were collected from descendant plants of each line. (**F**) Representative image of dithizone-stained grains showing zinc accumulation in wild-type and edited lines. (**G**) Grain zinc content (mg kg^−1^) in unpolished and polished rice. (**H**) Cadmium concentration (ppm) in grains of wild-type and edited lines, with the WHO permissible limit indicated. Pb and Cr levels were undetectable in grains.

Enhanced PEPC activity in the edited plants is expected to increase oxaloacetate production, thereby elevating the synthesis of malate and citrate. Citrate, in particular, chelates ferric ions to form ferric–citrate complexes that enhance Fe solubility and facilitate its mobilization in the rhizosphere (*16, 17*). Targeted metabolite profiling by GC–MS revealed substantial increase in organic acid levels in the edited plants, with citric acid elevated by 54% (151.4 ppm in edited versus 98.0 ppm in WT) and malic acid by 47% (67.2 ppm in edited versus 45.5 ppm in WT) relative to wild-type (Fig. 4C). Consistent with this mechanism, rhizospheric acidification assays showed that PEPC_R883G_ plants created a more acidic microenvironment around their roots than wild-type (Fig. 4D), a condition known to promote iron solubilization. Moreover, ferric chelate reductase (FCR) assays showed that roots of the edited plants were significantly more efficient at reducing Fe^3+^ to the bioavailable Fe^2+^ form than wild-type. On average, FCR activity was 64% higher in edited roots compared with WT (Fig. 4E). Together, these findings indicate that the edited plants possess an enhanced capacity for rhizospheric iron mobilization.

Because iron and zinc concentrations in rice grains are often positively correlated, and several transporters mediate the co-transport of both micronutrients (*18*), we next quantified zinc levels in the edited seeds. Dithizone staining produced markedly more intense coloration in grains from edited plants than in wild-type, indicating elevated zinc accumulation (Fig. 4F). Atomic absorption spectroscopy confirmed these observations: zinc content in unpolished grains increased by 40% (27.30 mg kg^−1^ in edited vs. 19.41 mg kg^−1^ in WT; Fig. 4G). Polished grains from edited plants similarly exhibited 42% higher zinc content than polished wild-type grains (21.71 vs. 15.27 mg kg^−1^) (Fig. 4G). In the best-performing line (line 63-4), zinc levels reached 28.29 mg kg^−1^ in unpolished grains and 22.51 mg kg^−1^ in polished grains. These metabolic shifts in PEPC_R883G_ plants likely increase the pool of bioavailable micronutrients entering the plant, ultimately supporting the substantial Fe and Zn enrichment observed in the seeds. Because this strategy enhances micronutrient acquisition through altered carbon flux and organic acid–mediated mobilization, rather than through overexpression of transporter or genes involved in iron accumulation (fig. S8), it is unlikely to broadly increase the uptake of toxic metals such as cadmium, chromium, or lead. Consistent with this expectation, targeted antinutrient profiling revealed no significant difference in cadmium accumulation between edited and wild-type plants, while chromium and lead levels were below the limits of detection (Fig. 4H).

Taken together, these data indicate that seeds from PEPC_R883G_ plants are substantially enriched in iron and zinc, levels consistent with long-standing biofortification targets proposed by the HarvestPlus program for improving nutrition in rice-dependent populations (*19*).

### Grains from prime-edited plants are biofortified with higher protein

Elevated PEPC activity in the edited plants is expected to increase oxaloacetate production— key precursors for the aspartate amino acid families. To assess whether this metabolic shift influences seed protein quality, we quantified total grain protein content (GPC). Among major cereals consumed as staple food, rice has one of the lowest intrinsic protein levels (*20*), and increasing GPC remains a longstanding goal for combating protein malnutrition in the developing world. An initial, non-destructive prediction using calibrated near-infrared (NIR) spectroscopy (*21*) indicated a marked 49.2% increase in GPC in unpolished edited seeds compared with wild-type (fig. S9). Consistent with this prediction, micro-Kjeldahl analysis confirmed that unpolished grains from the edited plants exhibited a 38.4% increase in GPC relative to wild-type (Fig. 5A-B). Edited unpolished grain contained, on average, 12.32% protein (range: 11.78–12.65%), compared with 8.9% in WT (range: 8.74 - 9.10%). Even after 1-minute polishing, edited grains retained an average of 10.1% protein (range: 9.28–10.61%), whereas polished WT grains contained only 6.7% (range: 6.48–7.02%), representing a 51% enhancement (Fig. 5A-B). This advance in the prime-edited lines is particularly notable, as successful breeding efforts for high-protein rice are scarce; to date, only two varieties have been released with ∼10% GPC (*22, 23*). Targeted metabolite analysis revealed a significant increase (34.25%) in oxaloacetate levels in the edited plants compared with wild-type (Fig. 4C). These results are consistent with a model in which elevated oxaloacetate availability enhances the supply of amino acid precursors to the developing grain. To test this possibility, we profiled seed amino acid composition using UPLC. As anticipated, edited seeds showed a pronounced increase in aspartic acid, accompanied by enrichment of nutritionally important amino acids, including lysine and the branched-chain amino acids leucine, isoleucine, and valine (Fig. 5C and fig. S10).

**Fig. 5:**
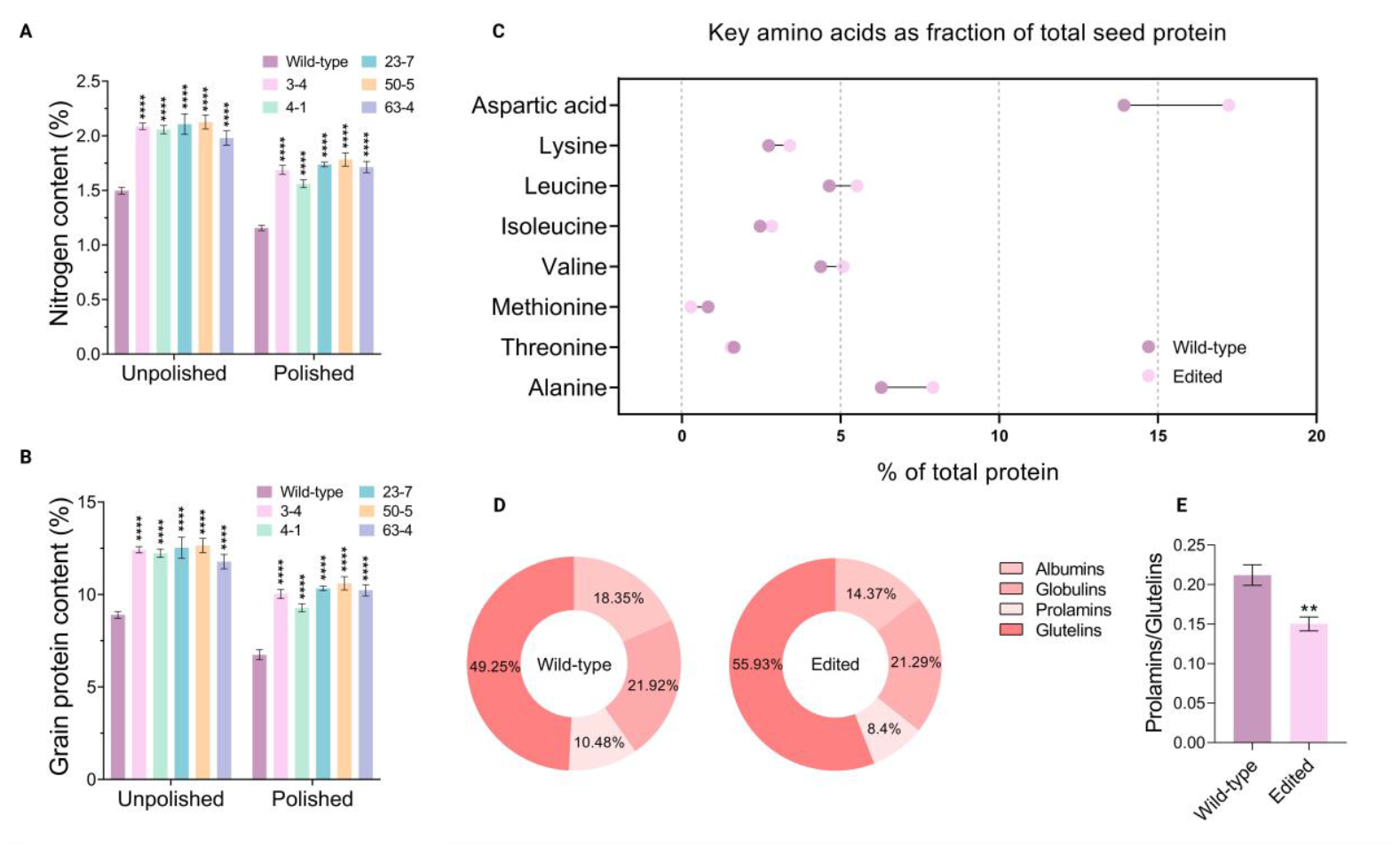
Enhanced protein accumulation in PEPC_R883G_ rice lines. (**A**) Total nitrogen content (%) in unpolished and polished grains of wild-type and edited lines analysed *via* micro Kjeldahl assay. (**B**) Grain protein content (%) in unpolished and polished rice of wild-type and edited lines. (**C**) Profile of selected amino acids (% of total seed protein); points represent mean values with connecting lines indicating the changes between wild and edited lines. (**D**) Relative distribution of major storage protein fractions (albumins, globulins, prolamins, and glutelins) in wild-type and edited grains. (**E**) Prolamins/Glutelins ratio in wild-type and edited grains. Data are presented as mean ± SD, and statistical significance relative to wild-type is indicated by asterisks (*P < 0.05, **P < 0.01, ***P < 0.001, ****P < 0.0001; ns, not significant).

Cereal grain storage proteins fall into four major classes—prolamins, glutelins, albumins, and globulins—distinguished by their solubility properties. Rice grain is nutritionally superior to barley, wheat, and maize because its dominant storage proteins are glutelins, which is more digestible and contains higher levels of lysine than prolamins (*24*). Protein fractionation revealed that the edited lines accumulated significantly higher levels of glutelins than WT (55.93% in edited versus 49.23% in WT; Fig. 5D and fig S11), aligning with long-standing breeding objectives to enhance glutelin levels in rice grain (*24*). Conversely, maintaining or reducing the prolamins-to-glutelins ratio is a key breeding objective for improving the nutritional value while preserving the eating quality of rice grain (*25*). Interestingly, edited grains contained 24.7% less prolamins than wild-type (Fig. 5D), resulting in a markedly lower prolamins-to-glutelins ratio (0.15 in edited lines versus 0.21 in WT) (Fig. 5E). Because changes in protein composition can influence grain texture and cooking related traits, we further assessed alkali spreading value (ASV) and starch content as indicators of grain quality. No significant differences in ASV or starch content were observed between wild-type and edited seeds (fig. S12 A-B).

Taken together, these results demonstrate that the PEPC_R883G_ modification can reprogram central carbon metabolism in ways that substantially enhance both the quantity and nutritional quality of grain protein.

### PEPC_R883G_ rice plants produce compact panicles with larger and heavier seeds

Beyond improved photosynthesis and enhanced nutritional value, the edited lines exhibited notable modifications in yield-related traits. Compared with wild-type, PEPC_R883G_ plants developed more compact panicles, characterized by reduced panicle length and markedly higher spikelet density (Fig. 6A-C). Architectural analysis revealed fewer primary branches but a significantly greater number of secondary branches, producing thicker, more compact panicles (Fig. 6D-G). Pedicel length was reduced by ∼46% relative to wild-type, contributing to the condensed panicle structure (Fig. 6H-I). Despite these architectural shifts, the total grain number per plant did not differ significantly between edited and wild-type plants (fig. S13). In addition, the edited plants showed early flowering and reduced plant height compared to WT plants (fig. S14).

**Fig. 6:**
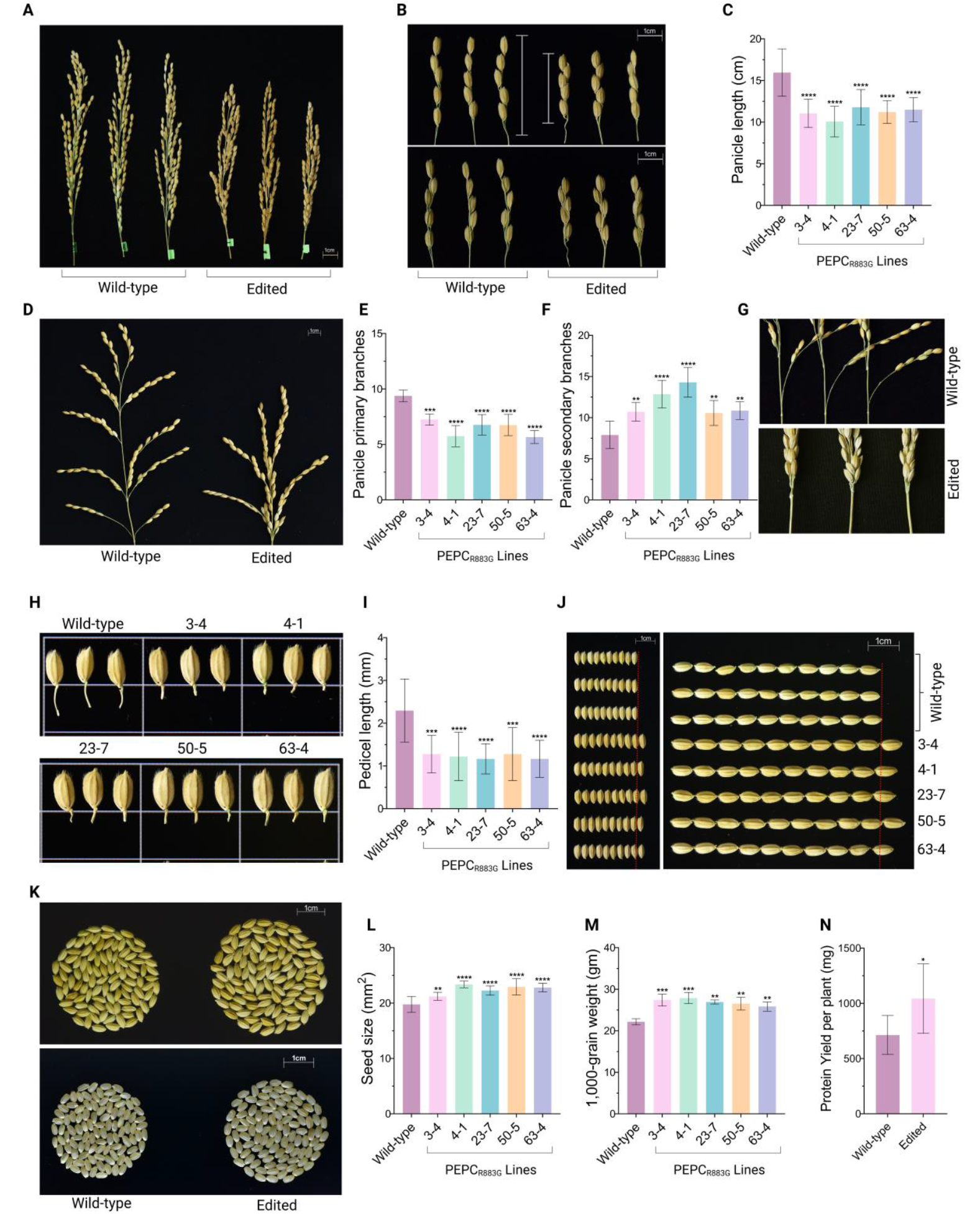
Effects of R883G substitution in PEPC on panicle architecture and grain yield–related traits in rice. (**A**) Representative images of mature panicles from wild-type and edited plants showing differences in overall panicle length. (**B**) Representative images illustrating the space occupied by five spikelets in wild-type and edited plants (scale bar = 1 cm). The upper panels show the adaxial side, and the lower panels show the abaxial side. (**C**) Panicle length in wild-type and independent edited lines. (**D**) Representative images showing overall panicle architecture and branching pattern in wild-type and edited plants. (**E**) Number of primary panicle branches. (**F**) Number of secondary panicle branches. (**G**) Enlarged views of spikelet arrangement at the base of the panicle in wild-type and edited lines. Panicles of PEPC_R883G_ plants exhibit a denser spikelet arrangement at the panicle base compared with the wild type. (**H**) Representative images of individual spikelets with pedicels. (**I**) Average pedicel length in wild-type and edited lines. (**J**) Representative images showing the comparison of grain length and width in wild-type and edited lines (scale bar = 1 cm). (**K**) Comparison of 100-grain size between wild-type and edited plants. (**L**) Average seed size (mm^2^) in wild-type and edited lines. (**M**) Thousand-grain weight (g) in wild-type and edited lines. (**N**) Protein yield per plant (mg), calculated by multiplying total grain weight per plant by the percentage of grain protein. Data are presented as mean ± SD, and statistical significance relative to wild-type is indicated by asterisks (*P < 0.05, **P < 0.01, ***P < 0.001, ****P < 0.0001).

Interestingly, we observed substantially altered seed morphology. Edited plants produced grains that were both longer and wider, resulting in a ∼14% increase in overall seed size (Fig. 6J-L and fig. S15). Consistent with these changes, we observed a significant increase (21%) in 1000-grain weight in PEPC_R883G_ plants (Fig. 6M). Interestingly, protein yield per plant was 46% higher in the edited lines than in wild-type plants (1,043 mg per edited plant versus 713.77 mg per WT plant; Fig. 6N).

Together, these findings indicate that enhancing PEPC activity through the R883G substitution not only boosts photosynthetic performance but also positively reshapes panicle architecture and seed development, leading to improvements in key yield-determining traits.

## Conclusion

In this study, we demonstrate a previously unexplored strategy for crop biofortification by enhancing grain zinc, iron, and protein (ZiP) content in rice through precise modulation of phosphoenolpyruvate carboxylase activity. By introducing a single, evolutionarily informed amino acid substitution into the endogenous C_3_ PEPC enzyme, we show that it is possible to simultaneously enhance photosynthetic performance, increase seed size, and enrich multiple essential nutritional components of rice grain. Notably, the concurrent enrichment of iron, zinc, and protein directly targets multiple, co-occurring nutritional deficiencies that underlie widespread malnutrition in populations reliant on cereal-based diets.

Unlike earlier iron and zinc biofortification efforts that relied largely on transgenic approaches (*26, 27*) and faced regulatory and public-acceptance barriers(*9*), the prime-editing strategy employed here generates transgene-free plants with single-base precision, substantially enhancing translational potential. Our findings establish PEPC as a powerful metabolic leverage point linking carbon assimilation, nutrient acquisition, and seed nutrient allocation in C_3_ rice. These findings provide a proof-of-concept for ZiP rice, illustrating how precise metabolic editing can couple enhanced photosynthesis with improved grain nutrition. Together, this work opens a new avenue for simultaneously improving photosynthesis and dietary quality in staple C_3_ cereals, with relevance for sustainable agriculture and human nutrition.

## Acknowledgements

We thank Dr. Pengcheng Wei (Anhui Agricultural University) for generously providing the enPPE2 plasmid. K.M. acknowledges funding support from the Ignite Life Science Foundation, Bangalore. S.P. acknowledges financial support from the Department of Science and Technology (DST), Government of India, through the INSPIRE programme. The authors acknowledge support from the Director, ICAR–CRRI, and the CRRI Central Laboratory Facility (CLF). CORE support from NABI to A.K.P. and the NASI Distinguished Honorary Professor Fellowship to S.K.D. are gratefully acknowledged. We also thank Dr. Sudipta Ray, Dr. Susmita Munda, and Mrs. Saloni Baskey for their assistance with various experiments and analyses.

## Author contributions

K.M. and M.J.B. conceived and designed the study and supervised the overall project. K.M., S.P., and T.H. planned the experiments. S.P. and T.H. performed the majority of the experiments and collected the data. S.P., T.H., K.M., T.A., R.K., A.M., and M.J.B. conducted data analysis. C.G., M.D., and S.P.A. assisted with cloning and tissue culture. A.T. assisted with protein-related work and screening of advanced mutant generations. D.P. assisted with rice protoplast transfection. S.K. constructed the pK-PE2 vector. D.B. assisted with gas-exchange measurements using IRGA. T.M. and K.M.K. performed molecular docking analyses. T.A. and A.M. carried out metabolite analysis and amino acid profiling. R.K. assisted with iron, zinc, and antinutrient profiling. S.B. conducted stomatal imaging. K.C. performed NIRS analysis and provided guidance on protein profiling. A.K.P., S.K.D., M.C., and M.K.K. provided advice on hypothesis development and experimental design. K.M., T.H., and S.P. wrote the manuscript. T.H., and S.P. prepared figures. K.M., M.J.B., K.C., A.K.P., S.K.D., T.A., R.K., M.C., and M.K.K. edited the manuscript. All authors contributed to the final version of the manuscript.

## Declaration of generative AI and AI-assisted technologies

The authors declare that GPT-5.2 was used to assist with text editing and proofreading. All generated content was subsequently reviewed, edited, and approved by the authors, who take full responsibility for the content of the publication.

## Data and materials availability statement

All data supporting the findings of this study are accessible within the article and its supplemental file. Amplicon sequencing data is available from NCBI (Bioproject ID, PRJNA1402686). Materials generated in this study, including the edited rice lines, are available upon reasonable request. Requests for mutant seed material must be submitted through the Director General, Indian Council of Agricultural Research (ICAR), New Delhi, and will be subject to institutional, national, and regulatory guidelines.

## List of supplementary materials

Materials and Methods

Figs. S1 to S15

Tables S1 and S3

Sequences S1 and S2

References

