## Supplementary material for "Prime editing of an evolutionarily informed PEPC residue enhances photosynthesis and grain nutrition in C_3_ rice": Supllemental file: Supplementary File_biorxiv - Copy.pdf

**The PDF file includes:**

- Materials and Methods
- Figures S1 to S15
- Tables S1 to S3
- Sequences S1 and S2

### Supplemental Materials and Methods

#### Multiple sequence alignment of C3 and C4 PEPC:

PEPC protein sequences from representative $C_3$ and $C_4$ plant species were retrieved from the NCBI /phytazome/ uniport database (accession numbers provided in Table S1). Multiple sequence alignment (MSA) was performed using Clustal Omega with default parameters to identify conserved and variable regions. The alignment was further inspected and manually curated in BioEdit. Particular attention was given to the residues encompassing the malate-binding site, and comparative analysis was carried out to detect amino acid substitutions or conserved motifs between $C_3$ and $C_4$ isoforms.

#### Protein-ligand docking analysis

The amino acid sequences of wild-type and mutant rice PEPC proteins were used to predict their three-dimensional structures using the AlphaFold Server (https://alphafoldserver.com/), followed by homology-based structural refinement using MODELLER. The modelled structures were then validated using the SAVES server (https://saves.mbi.ucla.edu/) with Ramachandran Plot analysis guiding the selection of protein models for further studies. Following validation, the ligand compound malate was retrieved from PubChem (https://pubchem.ncbi.nlm.nih.gov/) for docking studies. To investigate the interaction between malate and the PEPC, structure-based molecular docking was carried out using Schrödinger Maestro. PEPC structures were prepared using the Protein Preparation Wizard, which involved bond order assignment, addition of hydrogen atoms, optimization of hydrogen bonds, and energy minimization with the OPLS4 force field. The malate ligand was processed using LigPrep to generate 3D-optimized conformers with appropriate ionization states. Docking was performed using the Glide module in Standard Precision (SP) mode, with docking grids centered on predicted binding pockets identified using ClustalW (https://www.genome.jp/tools-bin/clustalw). The SP docking protocol provided detailed docking scores for the interaction between malate and the respective protein models.

#### Plant material and growth conditions

The rice cultivar *Kitaake* (*Oryza sativa* L.) was used as the plant material in the present study. Seeds were initially germinated on Murashige and Skoog (MS) medium, and 7-day-old seedlings were subsequently transferred to soil and grown under greenhouse conditions. Mature seeds were harvested, properly dried, and stored at room temperature for future use. Immature seeds were also collected as needed for callus induction throughout the study. For protoplast isolation, etiolated seedlings were used. Briefly, seeds were bisected, and the half containing the embryo was placed on MS medium. The seeds were germinated in darkness and allowed to grow for 10 days.

#### Construct generation for recombinant PEPC expression in *E.coil* cells

Total RNA was extracted from 100 mg of leaf tissue collected from one month old wild-type rice plants using NucleoSpin RNA Plant kit (MACHEREY-NAGEL). After spectrophotometric quantification, approximately 1 µg of total RNA was used for first-strand cDNA synthesis with the RevertAid First Strand cDNA Synthesis Kit (Thermo Scientific), employing oligo (dT) primers as per the manufacturer's instructions. The complete open reading frame (ORF) of OsPEPC (Accession no. OsKitaake08g131900) was subsequently PCR-amplified using gene-specific primers 574 and 575 (listed in Table S2). The resulting PCR product was cloned into the pUC19 vector (New England Biolabs-NEB). A mutant version of OsPEPC (C2647G in OsPEPC CDS) was also created through PCR based site-directed mutagenesis (SDM) using primer sets 576/579 and 580/575 (listed in Table S2). To express both wild-type and mutant forms of OsPEPC ($PEPC_{R883G}$) in a heterologous system, the respective coding sequences were subsequently cloned into the pRSET A expression vector (Thermo Scientific) at XhoI/Acc65I restriction sites. Sequences were verified by whole plasmid sequencing.

**Expression, purification, and enzyme assay of recombinant PEPCs**

The expression vectors harboring coding sequence of wild-type and $PEPC_{R883G}$ were transformed into *E. coli* Rosetta (DE3) pLysS cells (Sigma-Aldrich). Recombinant protein expression was induced with 1 mM IPTG at mid-log phase, once the culture reached the appropriate optical density. Cells were subsequently incubated at 30 °C for 12 hours to facilitate recombinant protein production. Successful expression of recombinant PEPC was verified by SDS–PAGE analysis.

For recombinant protein purification, Induced cells were sonicated in lysis buffer [20mM Tris (Sigma-Aldrich), 1mM EDTA (Sigma-Aldrich), 1% TritonX-100 (Sigma-Aldrich),10mM β-me (Sigma-Aldrich), and 2 mM PMSF (Sigma-Aldrich)] and the crude lysate was clarified by centrifugation at 12000 rpm for 10 min at 4°C. The resulting supernatant was used for urea solubilization, followed by Ni-NTA base affinity purification in presence of 6M urea. The eluted product was step dialysed to remove the urea in the refolding buffer (100 mM HEPES-NaOH (pH 7.5), 1 mM EDTA (Sigma-Aldrich), 10 mM $MgCl_2$ (Sigma-Aldrich), 1 mM DTT (Sigma-Aldrich), 10% glycerol (Sigma-Aldrich), and 2 mM PMSF). Buffer exchange was done using Amicon Ultra -4 centrifugal filter units (50kDa cutoff, Millipore). The purified protein was validated through western blotting using anti-His antibody (Thermo Scientific) and subsequently used for PEPC activity assay.
PEPC activity was measured using a modified spectrophotometric method (*28*). Purified protein was added to an assay buffer containing 100 mM HEPES-NaOH (pH 7.5), 10 mM $MgCl_2$, 1 mM $NaHCO_3$ (Sigma-Aldrich), 0.2 mM NADH (Sigma-Aldrich), 5 mM Glucose-6-phosphate (Sigma-Aldrich) and 12 U/ml malate dehydrogenase (MDH) (AMRESCO). The reaction was initiated by the addition of 4mM phosphoenolpyruvate (PEP) (Sigma-Aldrich) in the assay solution. Enzyme activity was monitored by measuring the decrease in absorbance at 340 nm ($A_{340}$) for 5 minutes, corresponding to the oxidation of NADH, using a spectrophotometer (Hitachi, USA). To evaluate the inhibitory effect of malate on PEPC activity, approximately 10 U of purified PEPC and $PEPC_{R883G}$ proteins were incubated in the presence (1 mM) or absence of malate (Sigma-Aldrich). The PEPC activity under these conditions was measured as described above.

**Prime editing Vector construction and pegRNA design:**

For targeting OsPEPC in rice, three different prime editing vectors—pK-PE2, pK-ePE2, and enpPE2 were used using either pegRNA or epegRNA.

*Making of pK-PE2 vector*
nCas9-L-M-MLV-RT cassette was synthesized in pBR322 vector background from BIOBASIC (Canada). To generate pK-PE2, the synthesized fragment was digested with BstBI (NEB) and EcoRI (NEB). The released fragment nCas9-L-M-MLV-RT was cloned into pRGEB32 (Addgene# 63142) vector background using the same sets of enzymes to generate pK-PE2. The final construct was confirmed through restriction digestion.

*Making of pK-ePE2 vector*

For making pK-ePE2 vector, two separate fragments- fragment 1 (flexible linker) and fragment 2 (NC-32aaL- M-MLV-RTΔRNAseH) were amplified from pK-PE2 and pH-ePPE (Addgene#183097), respectively, using Q5® High-Fidelity DNA Polymerase (NEB). The PCR reaction was carried out using 706/707 and 708/709 primer sets (Table S2). Subsequently, the two fragments were cloned into the pUC19 vector (NEB) using XbaI and BamHI restriction sites via three-way ligation to generate pUC19-NC-RT and confirmed through Sanger sequencing. Finally, the XbaI/BamHI-digested fragment from pUC19-NC-RT was cloned in the pK-PE2 vector to generate pK-ePE2. The final vector was confirmed through restriction digestion.

*enpPE2 vector*
The enhanced plant prime editing vector enpPE2 was used in this study (*29*).

The pegRNA for targeting OsPEPC was designed using PlantpegDesigner webtool (www.plantgenomeediting.net) (*30*). To enhance the stability of the 3′ end of pegRNA and prevent degradation from 3′- exonucleases, a RNA aptamer (tevopreQ1) was incorporated *via* an 8-nucleotide linker, resulting in an engineered pegRNA (epegRNA) (*11*). The designed pegRNA (Protospacer-Scaffold-RTT-PBS) and epegRNA (Protospacer-Scaffold-RTT-PBS-linker-tevopreQ1) were synthesized from Genscript (USA) and digested with BsaI restriction enzyme. Digested pegRNA and epegRNA expression cassettes were inserted into pK-PE2, pK-ePE2, and enpPE2 binary vectors to generate transformation ready constructs.

**Protoplast transfection**

Protoplasts were isolated from 10-day-old etiolated seedlings of the rice following the protocol established in our laboratory (*10*). For polyethylene glycol (PEG)-mediated transfection, 30 µg of plasmid DNA corresponding to each construct, along with pRGE-GFP (developed in our Lab - OsUbi:GFP) as a control, was introduced into the protoplasts. The transfected protoplasts were then incubated at 32 °C for 72 h with gentle agitation at 25 rpm.

**NGS sequencing and data analysis**

Next-generation sequencing (NGS) of PCR amplicons was employed for editing efficiency evaluation at the target sites. After 72 h of incubation, genomic DNA was extracted from transfected rice protoplasts using the NucleoSpin Plant II Mini kit for DNA (MACHEREY-NAGEL). Target sites were amplified with site-specific primers 898F and 899R (Table S2) containing Illumina adaptors using Q5® High-Fidelity DNA Polymerase (NEB). The purified PCR products were sequenced using Genewiz amplicon EZ sequencing service (Azenta Life Sciences, USA). Data analysis was performed by using the CRISPResso2 web tool (https://crispresso2.pinellolab.org/submission).

***Agrobacterium*-mediated rice transformation**

The best-performing construct enpPE2-epegRNA was introduced into *Agrobacterium tumefaciens* strain LBA4404-Vir2 (GOLDBIO) via electroporation, and positive colonies were confirmed by PCR analysis. Rice transformation was performed using immature seed-derived calli, following a previously published protocol (*31*). To evaluate editing efficiency in the early stage, hygromycin-resistant selected calli were used. Genomic DNA from selected calli was extracted using the NucleoSpin Plant DNA Isolation Kit (MACHEREY-NAGEL). The target region was subsequently amplified by PCR and subjected to deep sequencing as previously described (see section- NGS sequencing and data analysis). The transformed calli were subsequently regenerated as described earlier (*31*), and the regenerated plantlets were rooted, acclimatized and grown until maturity under greenhouse conditions.

**Screening of regenerated plants and identification of prime editor free mutants**

Genomic DNA was extracted from the leaves of $T_0$ putative edited plants using the cetyltrimethylammonium bromide (CTAB) method (*32*). The genomic region encompassing the target site was PCR amplified using gene-specific primers (1150F and 1151R) (listed in Table S2). The amplified products were subjected to Sanger sequencing, using primer 1357F (Table S2), to confirm the presence of desired edit. Seeds collected from edited $T_0$ plants were grown to obtain the $T_1$ plants. $T_1$ plants were screened for PEPC edit using same primer set and presence or absence of the Prime Editor (PE) components (Primer- 105 and 1386) and Cas9 transgenes (Primer- 100 and 900). Plants lacking both PE components and Cas9, yet retaining the desired edit at the target locus in both chromosomes (confirmed by Sanger sequencing), were identified and selected for further molecular and phenotypic analyses.

**Prediction and analysis of sgRNA-dependent off-targets**

Potential sgRNA-dependent off-target sites were identified using the CRISPR-GE webtool (https://skl.scau.edu.cn/) (*33*) and further validated through BLAST searches in the Rice Annotation Project Database (https://rice.uga.edu/) and Phytazome (https://phytozome-next.jgi.doe.gov/). Candidate off-target loci identified from these analyses (Table S3) were PCR-amplified using site-specific primers (listed in Table S2). The resulting amplicons were subsequently subjected to Sanger sequencing to determine the presence or absence of unintended edits.

**Quantitative RT PCR**

Expression of the PEPC in wild-type and the edited lines was determined via qRT-PCR. The actin and tubulin genes were used as an internal control for normalization of data. The qRT PCR was carried out using the QuantStudio 5 (Applied Biosystems, USA) with SYBR Green universal master mix (Applied Biosystems). The PCR cycling conditions were as follows: 95°C for 2 min, followed by 40 cycles of 95°C for 10 s and 58°C for 30 s. A melting curve analysis was routinely performed after 40 cycles to verify primer specificity and purity of amplicon. Similarly, expression of iron accumulation and transport related genes such as TOM1, NAS1, NAS2 were also determined via qRT-PCR. The primer sets used for expression study were listed in Table S2.

**Total protein extraction and PEPC activity assay**

Crude protein extracts were used for the PEPC activity assay as described earlier (*34*). Approximately 500 mg of leaf tissue was collected from both control and edited lines during late morning and immediately frozen in liquid nitrogen. Samples were ground to a fine powder with liquid nitrogen and homogenized in extraction buffer containing 100 mM HEPES-NaOH (pH 7.5), 1 mM EDTA, 10 mM $MgCl_2$, 1 mM DTT, 1% glycerol, 0.5% Triton X-100, 2 mM PMSF, and 0.1% protease inhibitor cocktail. The homogenate was centrifuged at 12,000 rpm for 30 minutes at 4°C, and the resulting supernatant was used for the PEPC activity assay. The PEPC activity assay was performed as described earlier (see section: Expression, purification, and enzyme assay of recombinant PEPCs).

**Photosynthetic pigment estimation**

Chlorophyll and carotenoid contents in the leaves of wild-type and edited plants were quantified following the method outlined by Arnon (1949) (*35*) , with minor modifications as described in Behera et al., 2023 (*3*). Approximately 25 mg of fully expanded leaf tissue was finely chopped and immersed in 5 ml of 80% (v/v) acetone and incubated at 4 °C in the dark for 48 hours to ensure thorough pigment extraction. After incubation, the supernatant was collected, and its absorbance was recorded at 480, 510, 645, and 663 nm using a UV–visible spectrophotometer (Hitachi, USA). The concentrations of total chlorophyll, chlorophyll a, chlorophyll b, and carotenoids were calculated using standard equations based on the absorbance values obtained from the acetone extract, and values were normalized to fresh tissue weight. The obtained values were expressed as mg $g^{-1}$ fresh weight of tissue.

**Measurements of photosynthetic gas exchange**

Gas exchange measurements were carried out on the fully expanded flag leaf at 50% flowering stage using a portable Infrared Gas Analyzer (IRGA) (LI-COR 6400 XT photosynthesis system; Lincoln, NE, USA). The measurements were taken under controlled conditions with a photosynthetically active radiation (PAR) of 1200 $\mu mol\ m^{-2}\ s^{-1}$, $CO_2$ concentration of 400 $\mu mol\ mol^{-1}$, chamber temperature maintained at 25 °C, and a flow rate of 500 $\mu mol\ s^{-1}$. Parameters such as net photosynthetic rate (A), transpiration rate (T), stomatal conductance (gs), light use efficiency, water use efficiency (WUE), and instantaneous water use efficiency (iWUE) were determined. In addition, $CO_2$ response (A/Ci) curve was generated following the manufacturer's guidelines to assess photosynthetic performance under varying intercellular $CO_2$ concentrations.

**Scanning electron microscopy for stomatal density**

Stomatal density was analysed in rice leaves using scanning electron microscopy (SEM, SNE-ALPHA) without chemical fixation. Fully expanded leaves were collected from both wild-type and edited plants of the same developmental stage and physiological condition. Leaf segments (~5 × 5 $mm^2$) were excised from the central region of the leaf blade, avoiding the midrib and margins, to ensure uniformity and reduce positional variability. Immediately after excision, leaf samples were gently air-dried in a desiccator at room temperature until complete surface moisture was removed. No chemical fixation or dehydration steps were employed, as the analysis was restricted exclusively to stomatal density measurements. The dried samples were mounted on aluminium SEM stubs with the abaxial leaf surface facing upward using double-sided carbon adhesive tape. Mounted samples were sputter-coated with a thin layer of gold (approximately 10–15 nm) to minimize charging during imaging. SEM observations were carried out using a scanning electron microscope operated at 20 kV and a working distance of approximately 6 mm. Images were captured at 1000X magnifications. Stomatal density was calculated as the number of stomata per unit leaf area (100µm × 100 µm). For each biological replicate, stomata were counted from multiple randomly selected fields of view, and the mean value was used for subsequent statistical analysis.

**Phenotypic evaluation of agronomic traits**

Agronomic traits of wild-type and edited rice plants were assessed following the Standard Evaluation System for Rice (SES) (*36*), which provides uniform criteria for trait measurement and comparison. In this evaluation, the heading date was recorded as the number of days from sowing until more than 50% of the plants exhibited panicle emergence above the flag leaf sheath. Plant height was measured at maturity as the vertical distance from the soil surface to the tip of the tallest panicle. To evaluate panicle architecture, the number of Primary and secondary branches per panicle, pedicel length, and the total spikelet count were manually recorded to capture variation in yield component. For grain morphology, ten well-filled grains were randomly sampled from both the basal and apical portion of the panicles from each plant to minimize sampling bias. Grain size parameters (length and width) were measured using *ImageJ* software based on high-resolution digital images. Grain weight was determined by weighing a batch of 200 grains, and the value was extrapolated to obtain the standard 1,000-grain weight.

**Perl's Prussian blue staining for the histochemical localization of iron**

In order to analyse the histochemical localization of iron, Perls' Prussian blue staining was performed following the method of M. Perls (1867) with minor modifications (*37*). Briefly, mature grains were manually dehusked and soaked in distilled water for 4 hr followed by sectioning and incubated in freshly prepared 2% potassium ferrocyanide (Sigma-Aldrich) and 2% HCl (MERCK) solution for 2 hr. Ferric ions released from the tissue reacted with potassium ferrocyanide to form an insoluble blue precipitate (ferric ferrocyanide), enabling detection of iron deposits.

**Dithizone staining for zinc detection**

Zinc distribution in rice seeds was assessed using a dithizone (DTZ)–based staining approach (*38*). Mature grains were manually dehusked, hydrated by soaking for 3–4 h, and longitudinally sectioned along the ventral crease prior to staining. A fresh dithizone staining solution was prepared by dissolving 1,5-diphenylthiocarbazone (0.5 mg $ml^{-1}$; Merck) in reagent-grade methanol. Seed sections were incubated in the staining solution for 20 min at room temperature in the dark. Zinc localization was detected by the appearance of a characteristic red–purple coloration corresponding to the formation of the Zn–dithizonate complex.

**Rice grain milling**

Mature rice grains from both wild-type and edited plant lines were harvested and air-dried to a uniform moisture content prior to milling. The brown rice (unpolished) samples were polished using a rice polisher (Zaccaria, Paz1dta) to remove the bran layer and obtain milled (polished) rice. Milling was performed for 1 minute under standardized conditions to ensure uniform removal of the outer layers across all samples. After milling, polished grains were immediately collected for downstream analyses. All samples were stored in airtight containers at room temperature for further biochemical or elemental analysis.

**Akali spreading assay**

The alkali spreading assay was performed to evaluate the gelatinization behaviour of rice grains following established method (*39*). For each set, six intact milled rice grains were immersed in 10 mL of 1.5% (w/v) KOH solution in a petri dish, ensuring that the grains did not touch each other. The dishes were covered and incubated at room temperature for 23 h. After incubation, the extent of grain spreading and disintegration was visually assessed.

**Estimation of iron, zinc, and antinutrient elements by atomic absorption spectrophotometry (AAS)**

Iron (Fe), zinc (Zn), and antinutrient elements including cadmium (Cd), lead (Pb), and chromium (Cr) concentrations in unpolished and polished rice grains from wild-type and edited lines were determined using a flame atomic absorption spectrophotometer (iCE™3500 AAS, Thermo Scientific, USA) equipped with element-specific hollow cathode lamps. Oven-dried rice seed samples (1000 mg) were finely ground using a mortar and pestle, and approximately 500 mg of powdered material was transferred into acid-washed microwave digestion tubes. Samples were digested with 5 mL of concentrated nitric acid ($HNO_3$) and 2 mL of hydrogen peroxide ($H_2O_2$, 30%) using a microwave digestion system (Ethos Easy, India) as described earlier (*40*). The digestion program ramped the temperature to 200 °C and held for 45 min. After cooling, digested samples were transferred to 25 mL volumetric flasks and diluted with ultrapure water. Blank digestions were carried out in parallel. Instrumental parameters were optimized prior to measurement. Iron and zinc measurements were carried out using an air–acetylene flame, while argon gas was used during the analysis of antinutrient elements to ensure stable atomization and improved sensitivity. The analytical wavelengths used were 248.3 nm for Fe, 213.9 nm for Zn, 228.8 nm for Cd, 217.0 nm for Pb, and 357.9 nm for Cr. Calibration curves were generated using certified standard solutions for iron, zinc, cadmium, lead, and chromium (Supelco, Sigma-Aldrich), and element concentrations were calculated from the corresponding absorbance values. All measurements were performed in triplicate, and concentrations were expressed on a dry weight basis.

**Rhizosphere acidification assay**

Rhizosphere acidification was examined using an agar-based pH indicator assay with bromocresol purple. Seeds were surface sterilized and germinated on Murashige and Skoog (MS) medium for five days under controlled environmental conditions. Uniform seedlings were then transferred to either iron-sufficient ($Fe^+$) or iron-deficient ($Fe^-$) MS medium and allowed to grow for an additional three days. After this treatment period, seedlings were placed on plates containing 1% (w/v) agar supplemented with 0.006% (w/v) bromocresol purple (Sigma-Aldrich) and 0.2 mM $CaSO_4$ (Sigma-Aldrich). The medium pH was adjusted to 6.5 using NaOH prior to solidification. Seedlings were incubated on these plates for 48 hr; plate images were recorded after 24 hr and 48 hr to assess rhizosphere acidification based on visible colour changes in the pH indicator.

**Ferric chelate reduction assay**

Ferric chelate reductase (FCR) activity was quantified using a ferrozine-based colorimetric assay. Seedlings were initially grown on Murashige and Skoog (MS) medium for seven days and subsequently transferred to iron-deficient ($Fe^-$) MS medium for three additional days. On the third day following transfer, approximately 100 mg roots were pooled into a single microcentrifuge tube, and immediately submerged in 700 µl of assay solution containing 0.1 mM Fe (III)-EDTA (Sigma-Aldrich) and 0.3 mM ferrozine (Sigma-Aldrich) prepared in distilled water. The reaction mixtures were incubated in darkness for 4 h to facilitate the reduction of $Fe^{3+}$ to $Fe^{2+}$ and subsequent formation of the $Fe^{2+}$ –ferrozine complex. Absorbance was then measured at 562 nm using a spectrophotometer. Ferric chelate reductase activity was calculated using the following equation:

$$\text{FCR activity}\ \left(\mu\text{M Fe}^{2+}\ \text{g}^{-1}\ \text{FW h}^{-1}\right) = \frac{\left(\frac{A}{28.6}\right) \times V}{\text{Root FW} \times T}$$

where $A$ denotes the absorbance at 562 nm, $V$ is the assay volume expressed in µl, *Root FW* represents the root fresh weight in grams, and $T$ is time interval in hours.

**Prediction of grain protein content using near-infrared reflectance spectroscopy (NIRS)**

Rice grain protein content was first predicted using near-infrared reflectance spectroscopy (NIRS) (DS2500F, FOSS). Approximately 5 g of brown rice was placed in a sample cup compatible with the NIRS instrument. Spectral data were collected over the near-infrared wavelength range 400–2500 nm at 2 nm intervals. Each sample was scanned in replicate, and the average spectrum was used for analysis. Protein content was predicted as percentage (%) on a dry weight basis.

**Estimation of Nitrogen by Kjeldahl method**

Total nitrogen content in unpolished and polished rice grains from both wild-type and edited lines was estimated using the Kjeldahl method (*41*). Briefly, 10 rice grains were mixed with 3 mL concentrated $H_2SO_4$ (Sigma-Aldrich) and 2 g of catalyst mixture (Sigma-Aldrich) ($K_2SO_4$ + $CuSO_4$ + Selenium) and heated until the dark solution turned clear, indicating complete conversion of organic nitrogen to ammonium sulphate. The digested samples were cooled, diluted with double-distilled water, and transferred to the distillation unit. During distillation, the ammonium ions were released as ammonia gas after the addition of 40% NaOH (Sigma-Aldrich) and captured in a receiver containing 4% boric acid solution, which facilitated quantitative trapping and colour change from violet to green. The trapped ammonia was then titrated against 0.01 N HCl (Sigma-Aldrich) using a mixed indicator (Methylene Blue and Methyl Red- Sigma-Aldrich) until the endpoint changed from green to violet. Nitrogen concentration was calculated using the formula:

$$\text{Nitrogen (\%)} = \frac{(V_b - V_s) \times \text{Strength of acid (N)} \times 1.4}{\text{Weight of sample } (W) \times 1000} \times 100$$

where $V_s$ = titration value of the sample, $V_b$ = titration value of the blank, $N$ = normality of HCl, $W$ = sample weight (g), and 1.4 is the conversion factor.

The final values were expressed as a percentage of dry weight, providing a reliable estimate of total nitrogen content and reflecting the plant's nitrogen assimilation efficiency. The Nitrogen content was multiplied by 5.95 to get the percentage of total grain protein content (*20*).

**Extraction and Fractionation of Rice Storage Proteins**

Rice storage proteins were sequentially extracted according to their solubility characteristics with minor modifications in the existing protocol (*20*). Polished rice grains from both wild-type and edited plants were ground into fine flour, and 2 g of flour was used for protein extraction. Rice flour (2 g) was defatted by adding 10 mL of n-hexane (Merck) and shaking vigorously for 60 min at room temperature. The mixture was centrifuged, and the supernatant was discarded. The defatting step was repeated twice. The defatted flour was air-dried at 45 °C to remove residual solvent. The dried, defatted rice flour was suspended in 10 mL of distilled water and stirred continuously at room temperature for 2 h. The mixture was centrifuged at 3000 × g for 30 min, and the supernatant was collected as the albumins fraction. The pellet obtained after albumins extraction was resuspended in 10 mL of 5% (w/v) NaCl (Sigma-Aldrich) solution and incubated with stirring at room temperature for 2 h. The sample was centrifuged at 3000 × g for 30 min, and the supernatant containing the globulins fraction was collected. The remaining pellet was extracted with 10 mL of 70% (v/v) ethanol (MERCK) and stirred for 2 h at room temperature. Following centrifugation at 3000 × g for 30 min, the supernatant was collected as the prolamins fraction. The final pellet was extracted with 10ml of 0.2 M sodium borate buffer (pH 10) supplemented with 0.5% (w/v) SDS (Sigma-Aldrich) and 0.6% (v/v) β-mercaptoethanol. The suspension was incubated at room temperature for 2 h with continuous mixing. After centrifugation, the supernatant was collected as the glutelins fraction. All extracted protein fractions were stored appropriately for downstream analysis. Protein quantification was carried out using Bradford reagent and visualised in SDS-PAGE.

**Targeted metabolite analysis**

Total metabolite profiling was conducted based on the methodology described previously with minor modification (*42*). Approximately 50 mg each of rice leaf was homogenized in 1 mL of methanol (MERCK), containing 40 µL of adonitol (Sigma-Aldrich) (2 mg/mL in water) as an internal standard. The homogenate was vortexed thoroughly and incubated at 70°C for 15 minutes with continuous shaking at 950 rpm. Subsequently, an equal volume of water and 500 µl of chloroform were added, and the mixture was centrifuged at 2200 × g for 15 minutes. A 250 µL aliquot of the methanol-water supernatant was collected, evaporated to dryness using a speed vacuum concentrator, and stored at −80°C until further use.
For derivatization, the dried extracts were resuspended in 40 µL of methoxyamine hydrochloride solution (Sigma-Aldrich) (20 mg/mL in pyridine) and incubated at 37°C for 2 hours at 600 rpm. Trimethylsilylation was carried out by adding 50 µL of N, O-bis (trimethylsilyl) trifluoroacetamide (BSTFA) (Sigma-Aldrich), followed by a further incubation at 37°C for 30 minutes. Finally, 1 µL of the derivatized sample was injected into a GC-MSMS system (Trace GC Ultra-TSQ 9000, Thermo Scientific USA) with Triplus RSH autosampler for metabolite detection and quantification in split less mode. The injector temperature was kept at 250ºC, and both MS transfer line and ion source were kept at 280ºC. Chromatographic separation was achieved using an SH-5MS capillary column (30 m × 0.25 mm i.d.; Shimadzu Corp., Japan). Helium (99.999% purity) was used as the carrier gas at a constant flow rate of 1.0 mL $min^{-1}$, and argon (99.999% purity) served as the collision gas. The oven temperature program was as follows: initial temperature of 70 °C held for 2 min, ramped to 100 °C at 5 °C $min^{-1}$ with a 2 min hold, followed by an increase to 280 °C at 15 °C $min^{-1}$ and a final hold for 2 min. Quantitative analysis was performed in selected reaction monitoring (SRM) using the characteristic ions and retention times (RT) of the detected metabolites as provided in table below (*43*). Prior to statistical analysis, the peak area of each metabolite in each sample was normalized to that of the internal standard (adonitol). The mass transitions used for quantification of the analytes are provided in the table.

| Compound | RT | Mass | Product Mass | Collision Energy |
|---|---|---|---|---|
| Oxaloacetic acid | 11.69 | 220 | 73.1 | 18 |
| Succinic acid | 14.58 | 147.1 | 73.1 | 14 |
| Fumaric acid | 14.97 | 245.1 | 73.1 | 18 |
| Malic acid | 16.54 | 233.1 | 73.1 | 10 |
| Ketoglutaric acid | 17.33 | 198.1 | 73.1 | 12 |
| Adonitol | 18.57 | 217 | 73.1 | 18 |
| Citric acid | 19.25 | 273.1 | 73.1 | 18 |

### Amino acid profiling

Amino acid composition was determined by following method described earlier (*44*, *45*). Briefly, approximately 5 mg of sample was taken in a 10 ml glass tube and hydrolysed with 6N hydrochloric acid at 110°C under anaerobic condition for 16 hours. The hydrolysed samples were neutralized with 6N NaOH and then derivatized using amino acid derivatization kit (Tag Ultra derivatization kit Waters part#186003836, Waters Corp, USA). 1 µl of derivatized sample was injected into the UPLC system with fluorescence detector (Waters Acquity UPLC H Class). The mobile phase and gradients programs were used as per the manual provided by the manufacturer (Acquity UPLC H-Class and H-Class Bio Amino acid Analysis System guide, 2012). The amino acids were quantified by comparing with the retention time and peak areas of the amino acid standards (Part No. WAT088122, Waters Corp. USA).

### Statistical analysis

All statistical figures were generated, and data analyses were performed using GraphPad Prism and BioRender software. Comparisons between wild-type and edited lines were conducted using One-way ANOVA and *t*-test wherever applicable. Statistical parameters, measures of central tendency (mean), and variability (± SD), as well as levels of statistical significance, are reported in the corresponding figures and figure legends. No data were excluded from the analysis, and plant materials were randomly selected from a sufficiently large pool to minimize sampling bias.

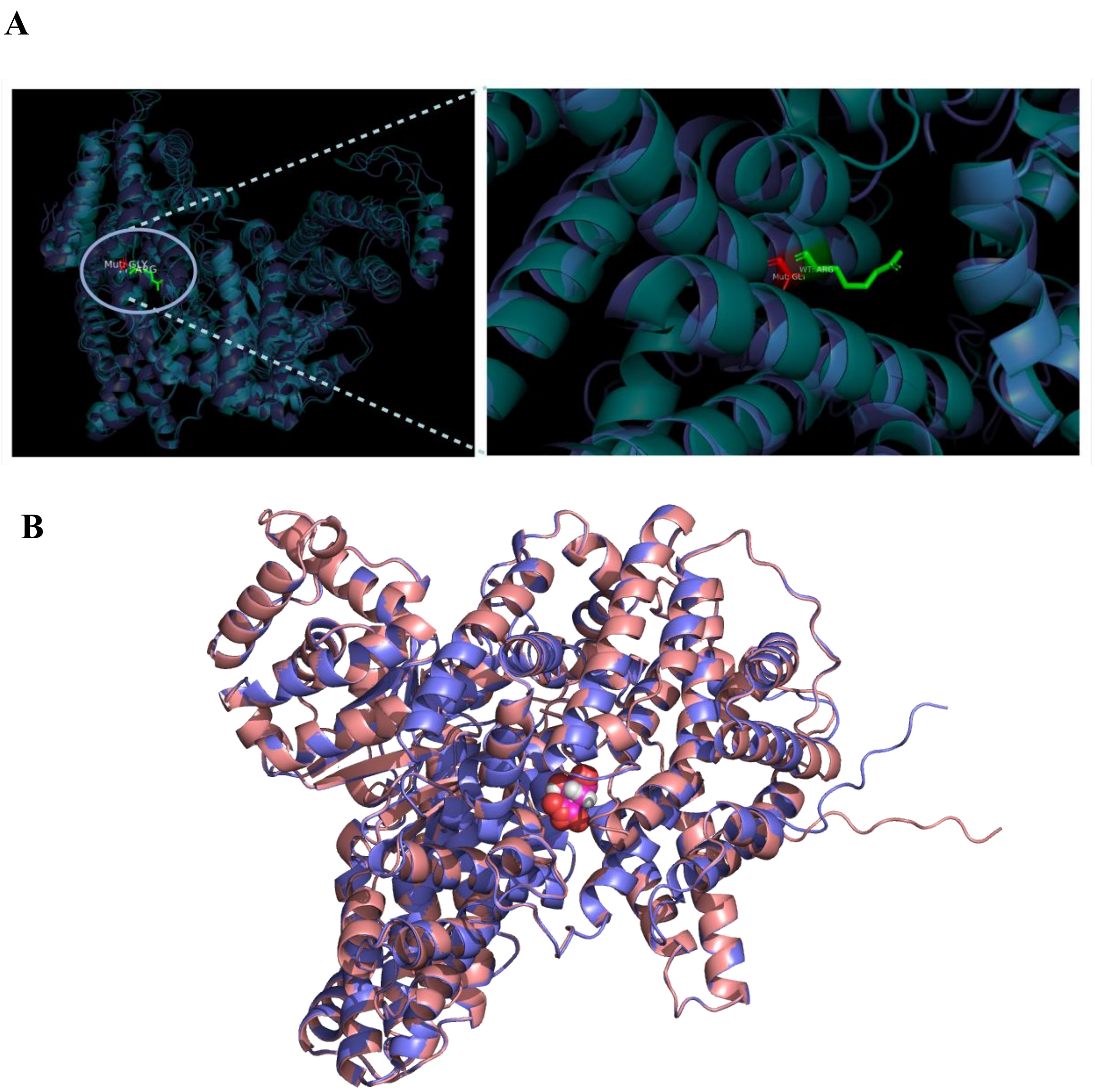

**Fig. S1. Structural representation of the PEPC protein showing Arg883⟶Gly883 substitution.**

(**A**) Three-dimensional structures of the PEPC protein depicting the wild-type and edited forms are shown in ribbon representation. The mutated residue is highlighted at the indicated position, where arginine (Arg883) in the wild-type protein is substituted by glycine (Gly). The left panel shows the overall protein structure with the mutation site highlighted, while the right panel presents a magnified view of the local structural environment surrounding the substituted residue, indicating the precise position of the Arg→Gly change. (**B**) Superimposed structure of the wild-type and edited PEPC, demonstrating that the amino acid substitution does not cause major alterations in the overall protein fold. Wild-type and edited proteins are shown in red–salmon and blue–slate, respectively.

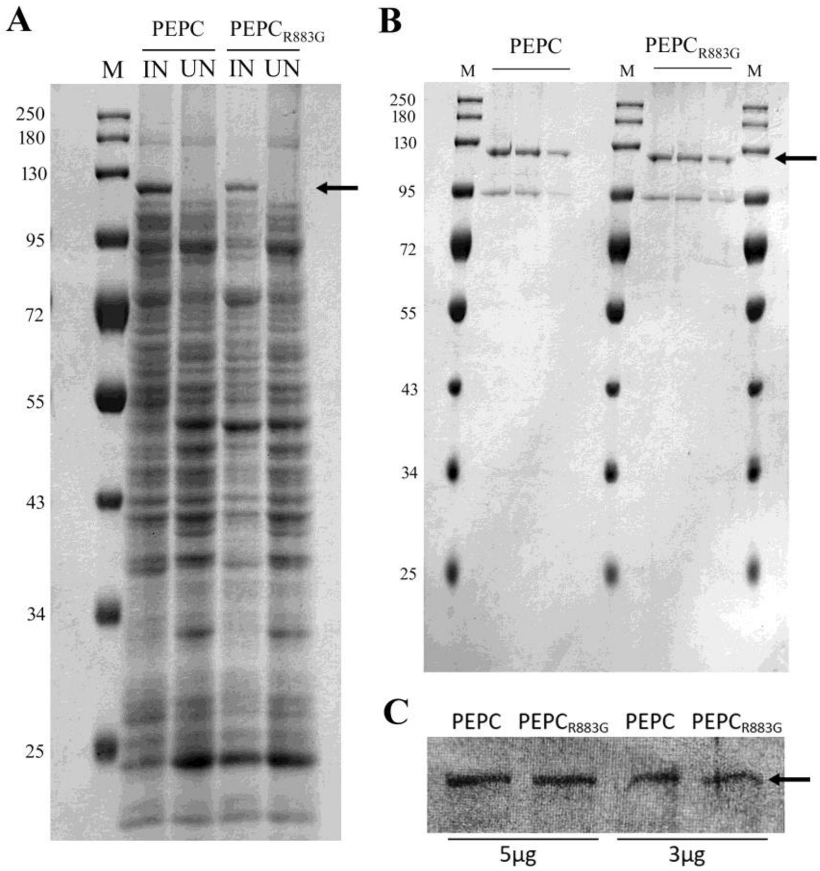

**Fig. S2: Expression and detection of PEPC and PEPC$_{R883G}$ protein by SDS–PAGE and immunoblot analysis.**

(**A**) Coomassie Brilliant Blue–stained SDS–PAGE showing the expression of recombinant PEPC (wild-type) and PEPC$_{R883G}$ (mutant) upon induction in *E. coli* Rosetta (DE3) pLysS. Lane M, protein molecular weight marker; IN, Induced; Un, Uninduced. (**B**) SDS–PAGE of affinity-purified PEPC and PEPC$_{R883G}$ protein, showing bands at the expected molecular weight (128 kDa). (**C**) Immunoblot analysis confirming the PEPC and PEPC$_{R883G}$ protein in the corresponding samples.

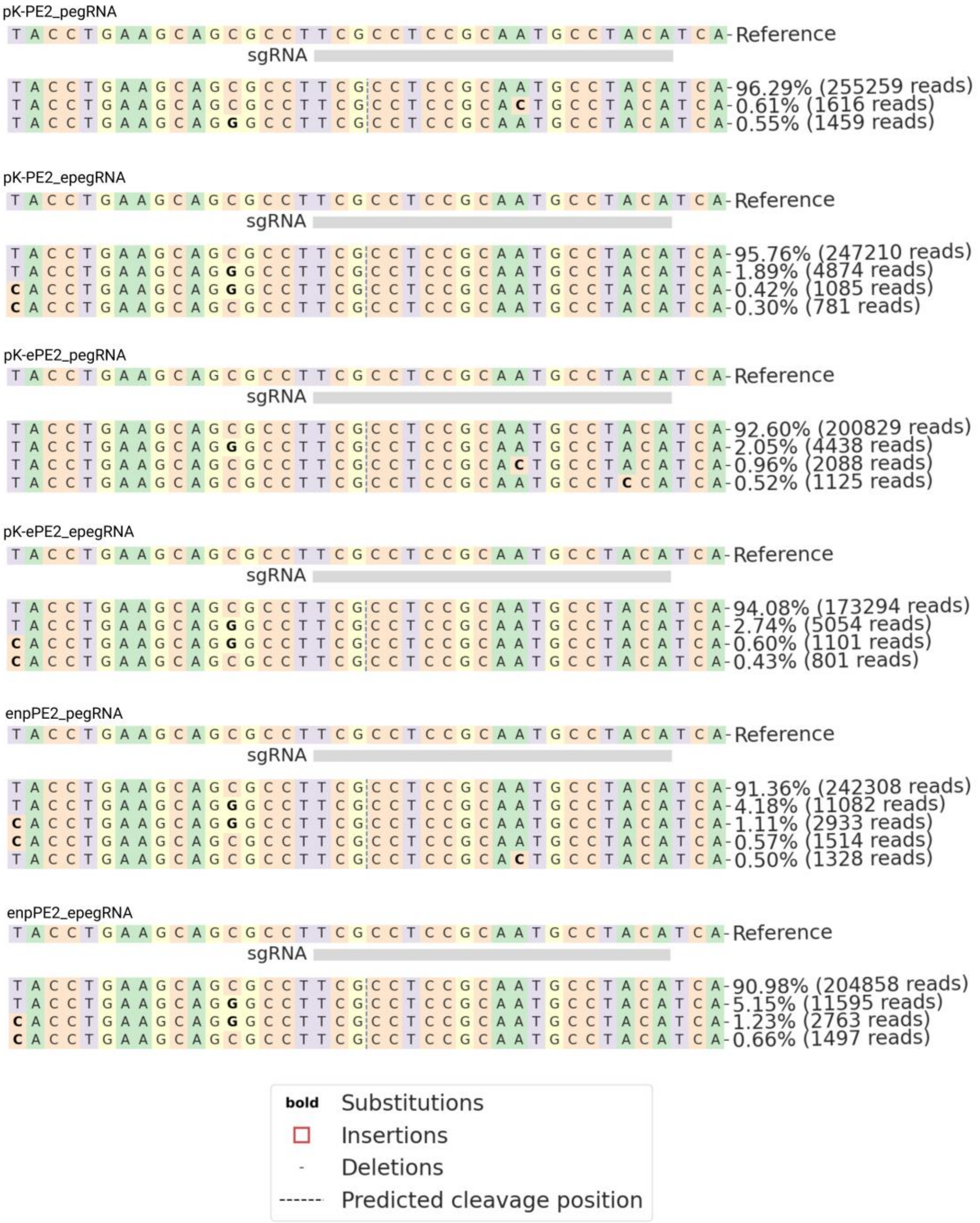

**Fig. S3: Prime-editing (C-to-G) efficiencies with different prime editing constructs using pegRNA and epegRNA.**

Screenshot from CRISPResso2 analysis depicting prime-editing outcomes for pK-PE2_pegRNA, pK-ePE2_pegRNA, pK-ePE2_epegRNA, enpPE2_pegRNA, and enpPE2_epegRNA constructs. The reference sequence and sgRNA target region are shown at the top of each panel. Aligned amplicon reads are displayed below, with allele frequencies (%) and corresponding read counts indicated on the right.

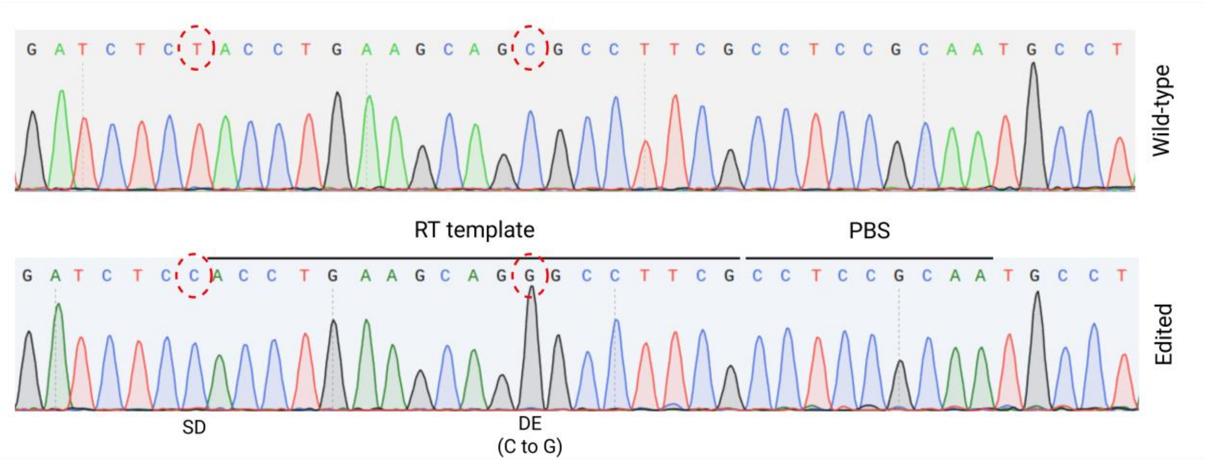

**Fig. S4**: **Representative Sanger sequencing chromatogram showing a byproduct derived from the epegRNA scaffold.**

Sanger chromatograms showing scaffold-derived (SD) byproduct along with the desired editing (DE) in the target locus. The red dashed circles indicate the edited nucleotide position.

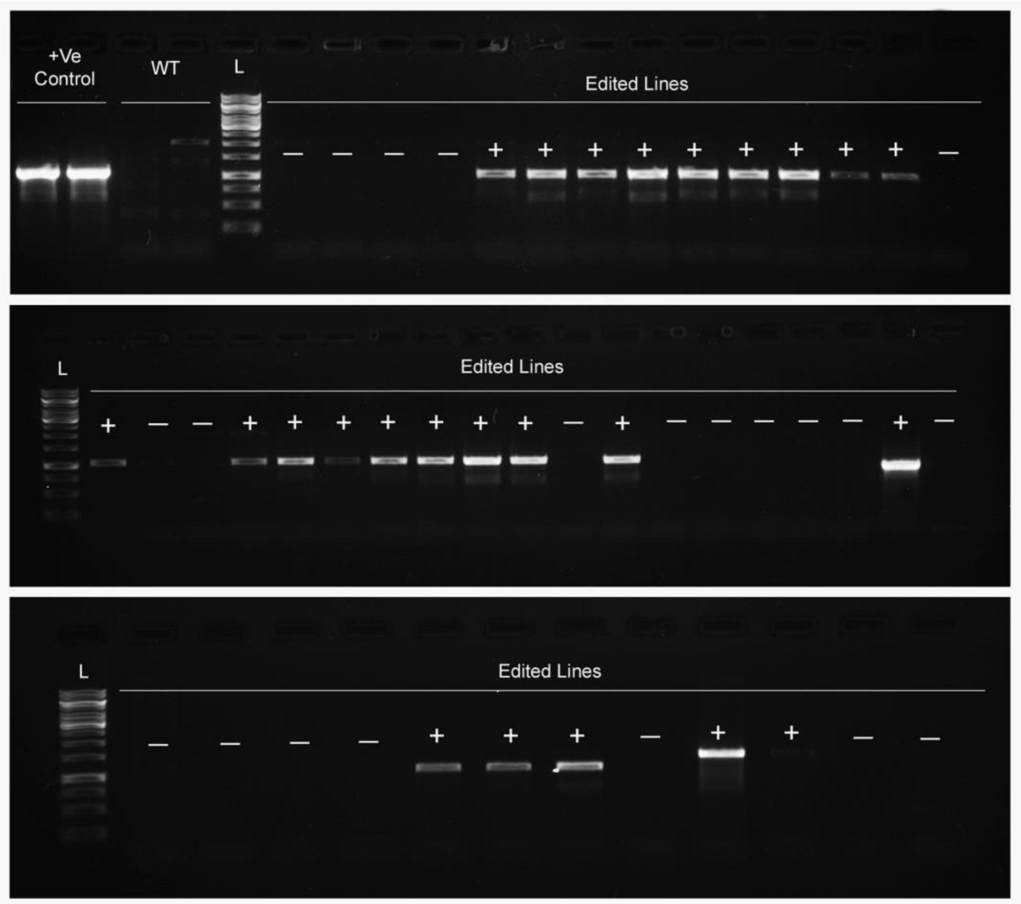

**Fig. S5: PCR-based genotyping to identify PE-free edited (PEPC$_{R883G}$) rice lines.**
Agarose gel electrophoresis showing amplification of a 1058-bp reverse transcriptase (RT) fragment from wild-type (WT), positive control (+ve; enpPE2-epegRNA plasmid), and representative edited lines. Lane L denotes the DNA ladder. Presence (+) or absence (−) of the expected amplicon indicates retention or loss, respectively, of prime-editor components in independent lines.

**A**

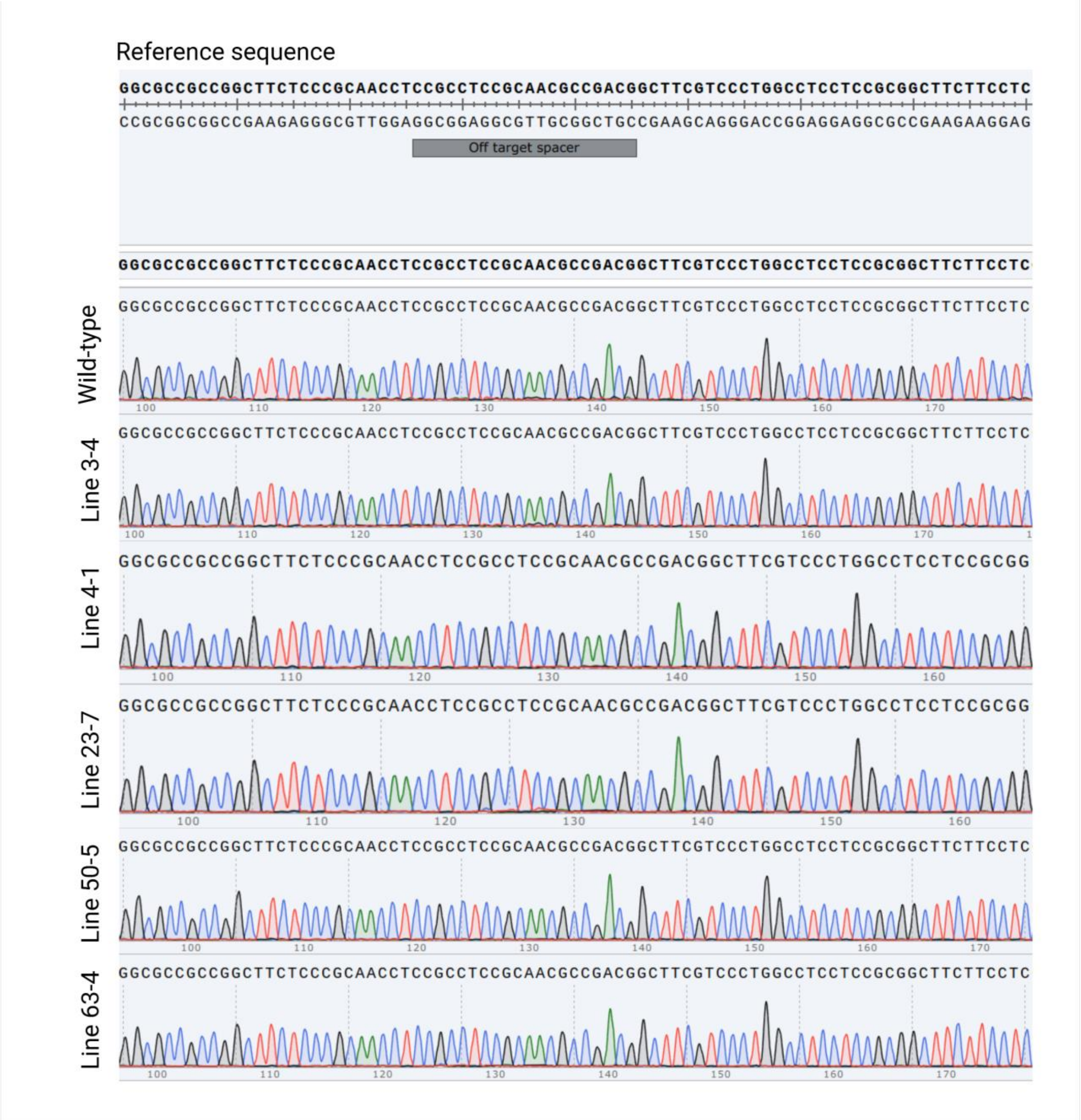

Reference sequence
GGCGCCGCCGGCTTCTCCCGCAACCTCCGCCTCCGCAACGCCGACGGCTTCGTCCCTGGCCTCCTCCGCGGCTTCTTCCTC
CCGCGGCGGCCGAAGAGGGCGTTGGAGGCGGAGGCGTTGCGGCTGCCGAAGCAGGGACCGGAGGAGGCGCCGAAGAAGGAG
Off target spacer
Wild-type
Line 3-4
Line 4-1
Line 23-7
Line 50-5
Line 63-4

**B**

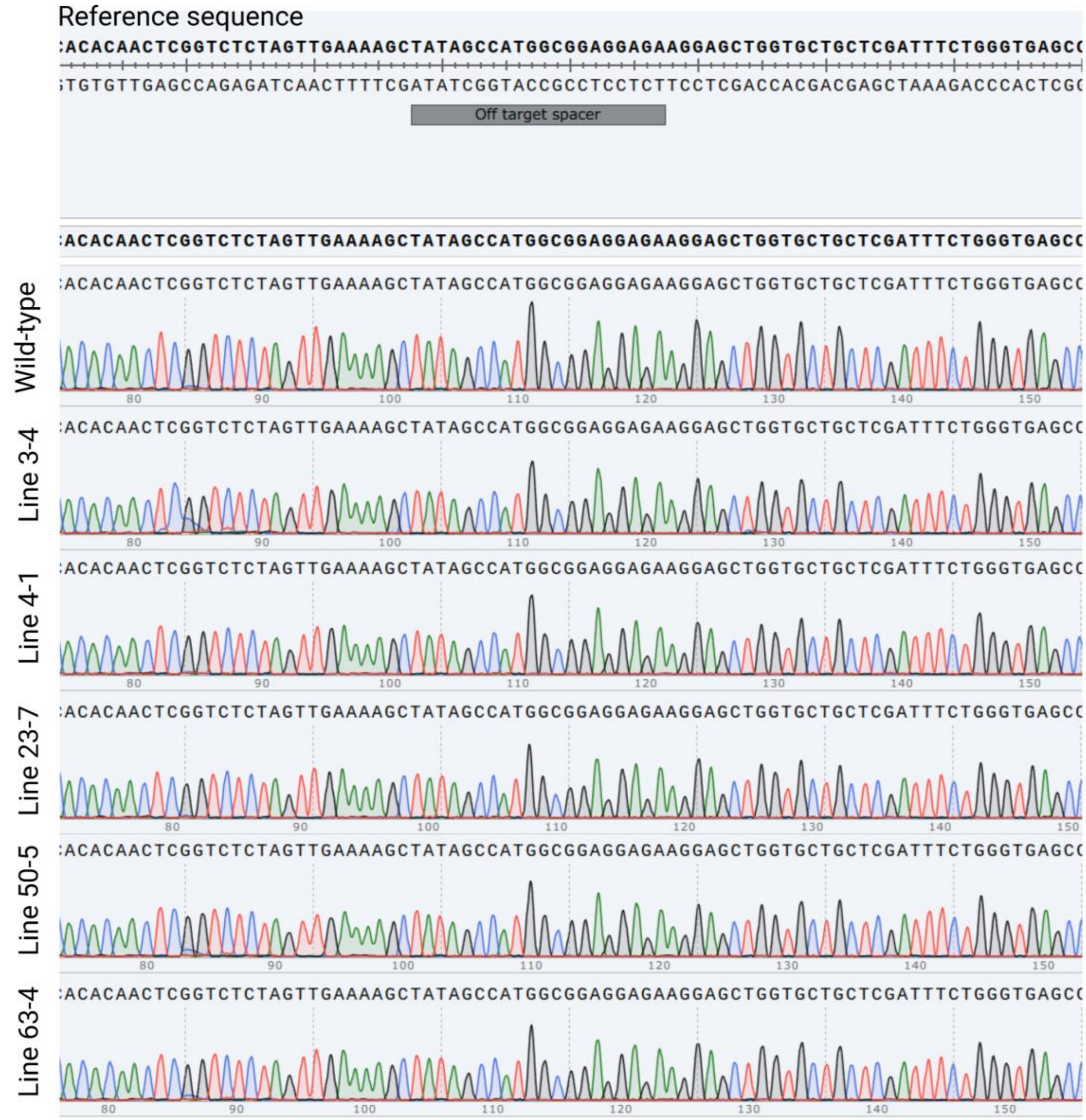
Reference sequence
:ACACAACTCGGTCTCTAGTTGAAAAGCTATAGCCATGGCGGAGGAGAAGGAGCTGGTGCTGCTCGATTTCTGGGTGAGCC
;TGTGTTGAGCCAGAGATCAACTTTTCGATATCGGTACCGCCTCCTCTTCCTCGACCACGACGAGCTAAAGACCCACTCGC
Off target spacer
Wild-type
Line 3-4
Line 4-1
Line 23-7
Line 50-5
Line 63-4

**C**

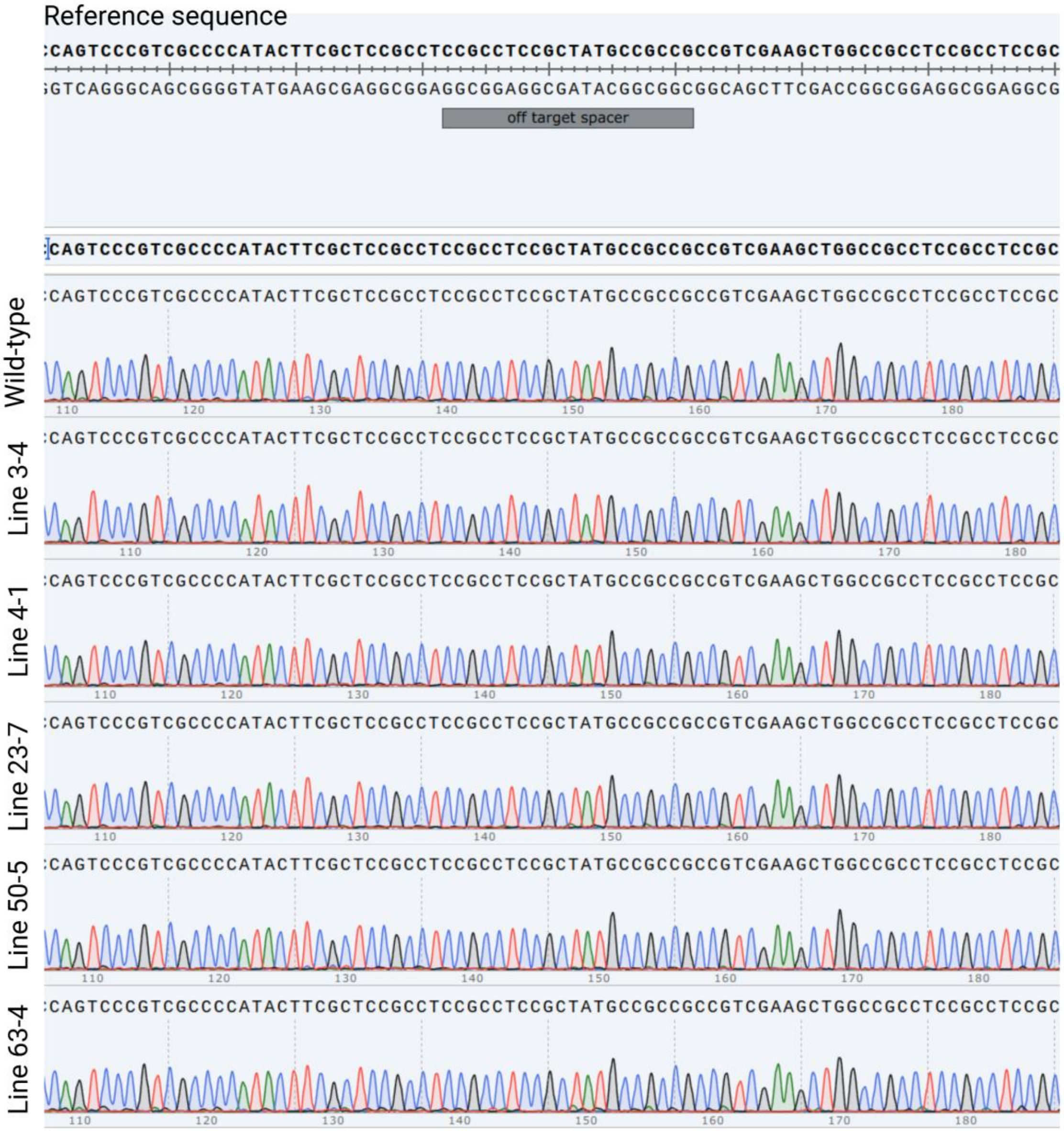

**Fig. S6: Prediction of off-target sites and analysis through Sanger sequencing in wild-type and edited lines.**

(**A**) Sanger sequencing chromatogram of predicted off-target site 1 (OsKitaake12g123300 Chr12_16070546...16072093). (**B**) Sanger sequencing chromatogram of predicted off-target site 2 (OsKitaake03g365700_Chr3_33899807...33900915) (**C**) Sanger sequencing chromatogram of predicted off-target site 3 (OsKitaake08g112366_Chr8_14286916...14290701). The off-target spacer sequence is indicated at the top of chromatograms. Representative chromatograms from wild-type and multiple independently edited lines show identical peak patterns, with no nucleotide substitutions or mixed peaks detected at the predicted off-target loci, confirming the absence of off-target mutations in the edited lines.

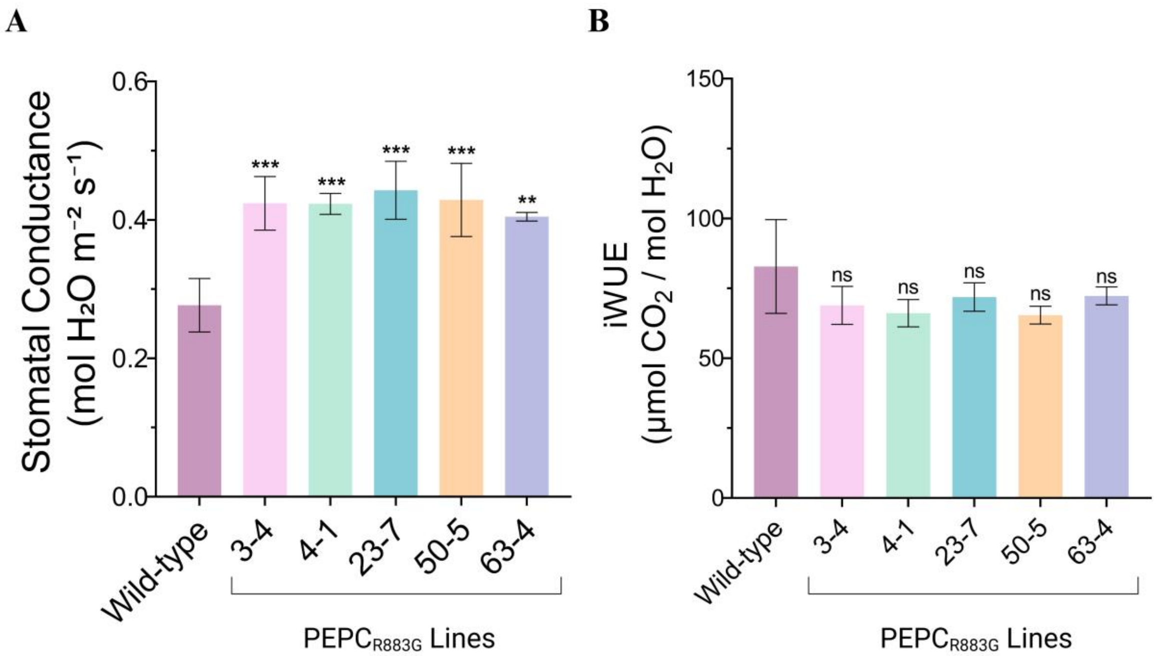

**Fig. S7: Stomatal conductance and intrinsic water use efficiency (iWUE) of wild-type and edited rice lines.**

(**A**) Stomatal conductance (mol $H_2O$ $m^{-2}$ s) was derived from transpiration rate and vapor pressure deficit. (**B**) iWUE was calculated as the ratio of net $CO_2$ assimilation rate to stomatal conductance and expressed as μmol $CO_2$ $mol^{-1}$ $H_2O$. No statistically significant differences in iWUE were observed between the wild-type and edited lines (ns, not significant). Bars represent mean ± SD of biological replicates. Asterisks indicate statistically significant differences relative to the wild type (**$p < 0.01$; ***$p < 0.001$).

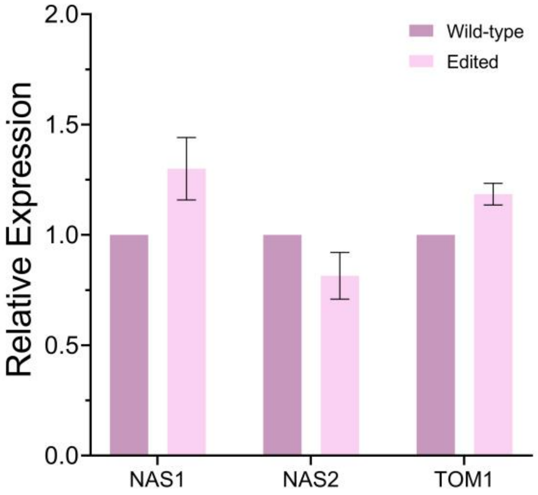

**Fig. S8: Relative expression levels of *NAS1*, *NAS2*, and *TOM1* genes in wild-type and edited rice lines.**

Gene expression was quantified by qRT-PCR and normalized to the internal reference gene (Actin), with the wild-type expression level set to 1. Bars represent mean relative expression values, and error bars indicate mean ± SD from biological replicates.

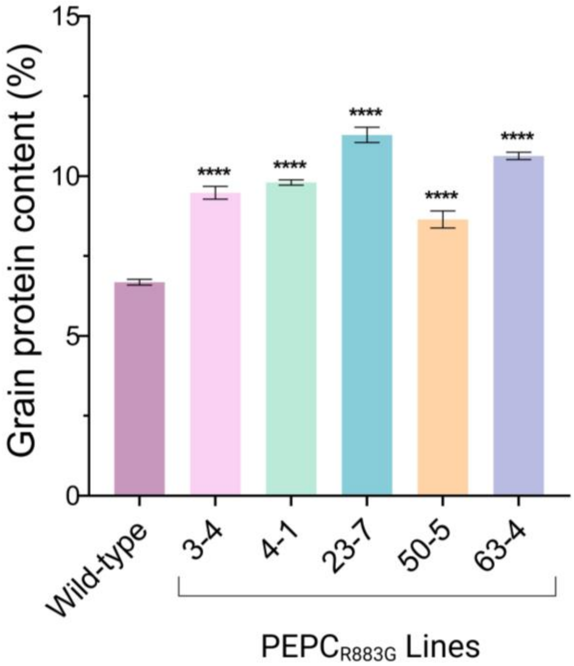

**Fig. S9: NIRS based prediction of Grain protein content (%) in wild-type and edited rice lines.**
Bars represent the mean grain protein content in wild-type and edited rice lines. Edited lines show significantly higher protein content compared with the wild type. Error bars indicate mean ± SD. **** denotes $P < 0.0001$.

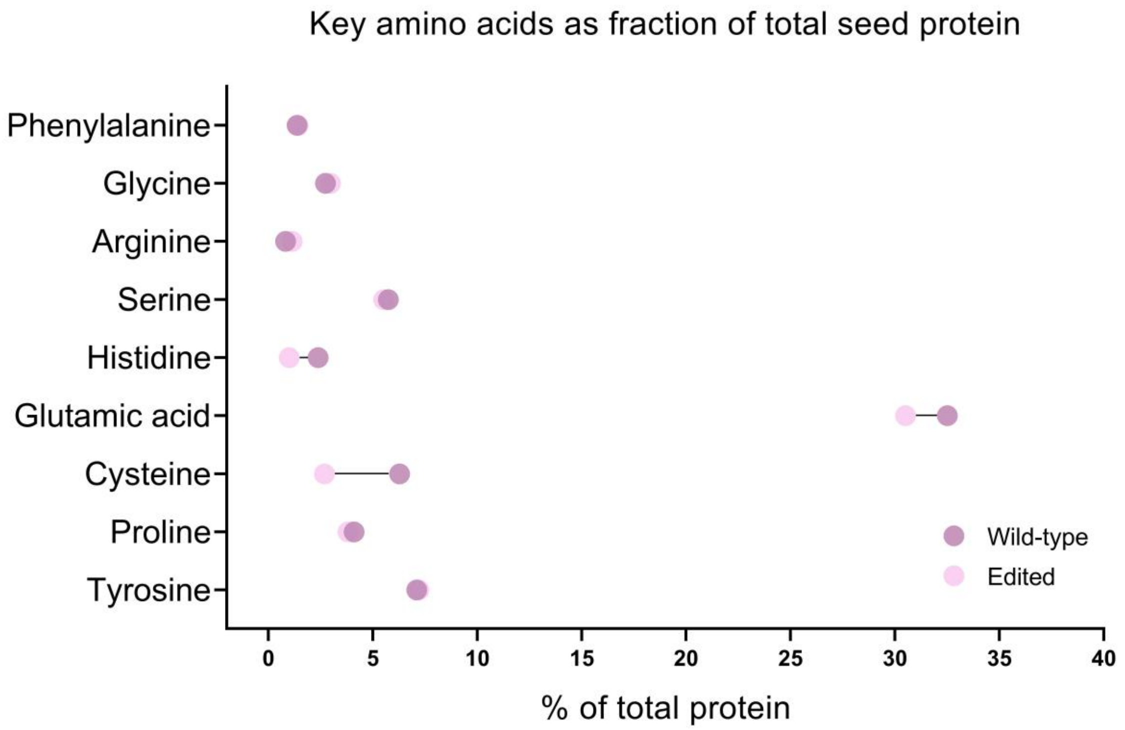

**Fig. S10: Comparison of selected amino acid composition between wild-type and edited polished grains**.
Points represent mean values with connecting lines indicating the changes in amino acid composition (% of total seed protein) between wild and edited lines.

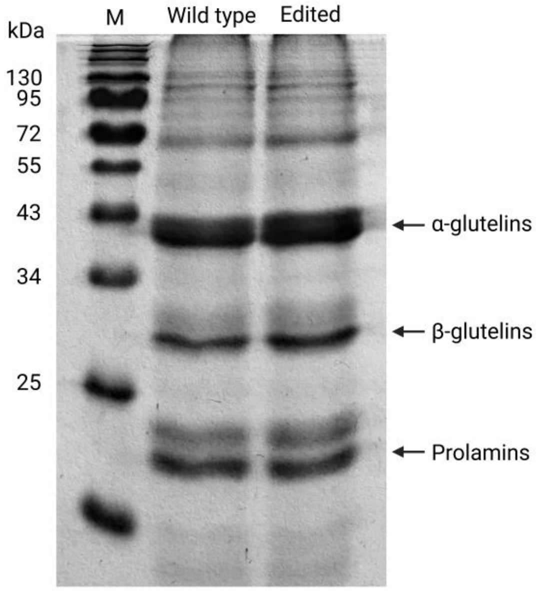

**Fig. S11: SDS–PAGE analysis of seed storage proteins in wild-type and edited.**
Different grain protein fractions were separated in SDS–PAGE and visualized by staining. Polished grains were used in fractionation. Lane M indicates the protein molecular weight marker. Distinct bands corresponding to α-glutelins, β-glutelins, and prolamins are indicated.

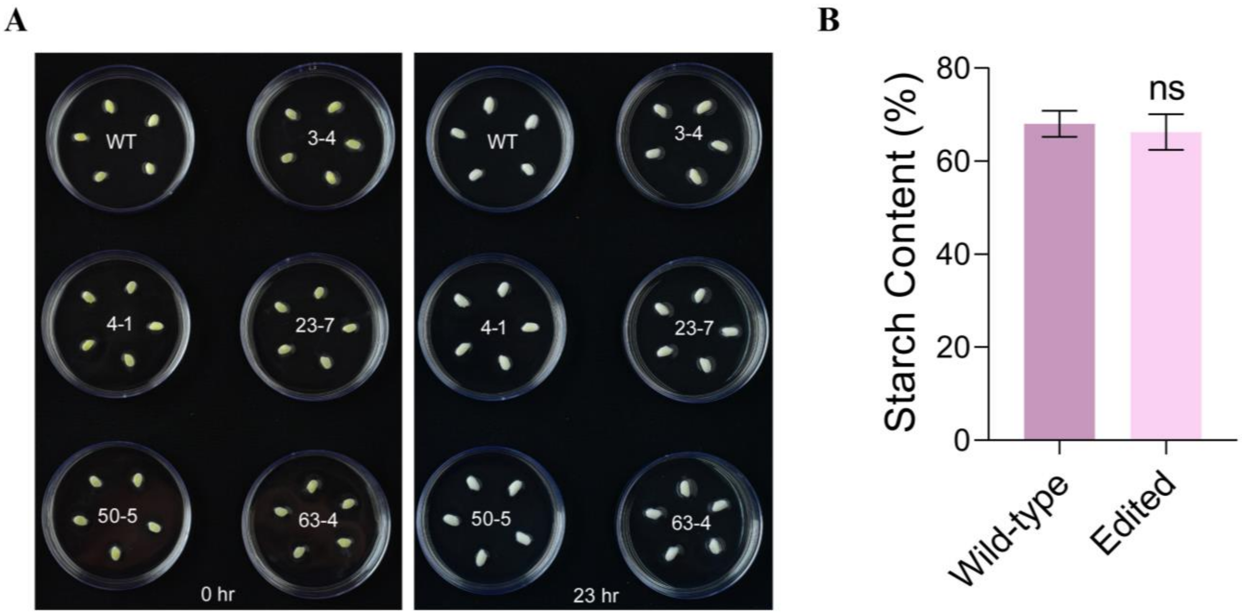

**Fig. S12: Alkali spreading value and starch content of seeds**
(**A**) Alkali spreading value (ASV) assay of wild-type (WT) and edited polished rice seeds. Representative images showing seeds of WT and edited lines (L-3, L-4, L-23, L-50, and L-63) at 0 h and 23 h after incubation in 1.5% KOH solution. (**B**) Total starch content was quantified in mature rice grains of wild-type and edited lines. Bars represent mean ± SD from biological replicates. No significant difference in starch content was observed between the wild type and edited lines (ns, non-significant).

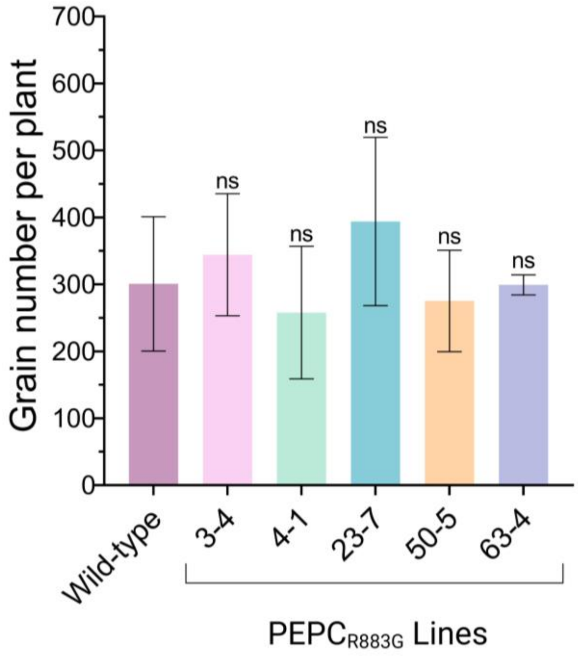

**Fig. S13: Grain number per plant**

Bars represent the mean grain number per plant, and error bars indicate mean ± SD from biological replicates. Data from eight homozygous PE-free plants from each line were calculated. ns, not statistically significant.

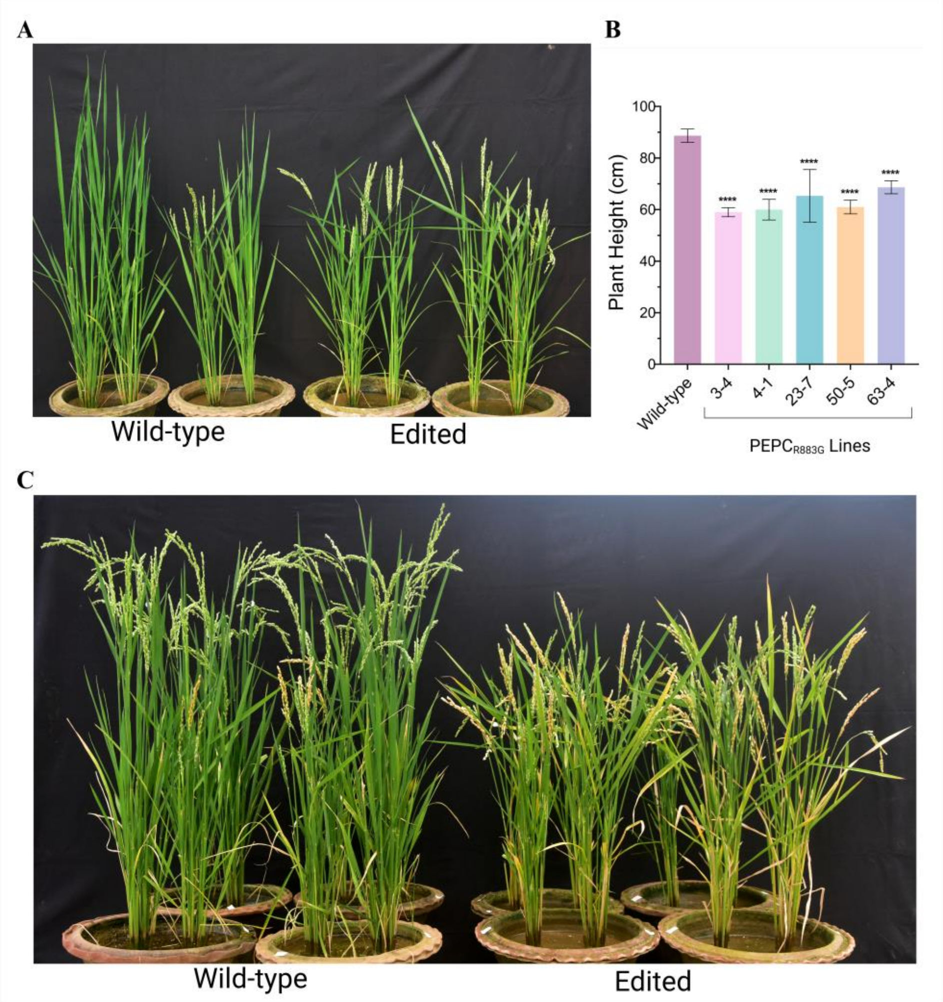

**Fig. S14: Phenotypic appearance of wild-type and edited rice lines.**
(**A**) Representative image showing the early flowering (16 days) phenotype in edited plants compared to that of wild-type plants grown under identical conditions. (**B**) Plant height (cm) in wild-type and edited lines. Bars represent mean values ± SD. Asterisks indicate statistically significant differences compared with the wild type (****, $P < 0.0001$). (**C**) Representative image of plants illustrating differences in plant height in wild-type and edited lines.

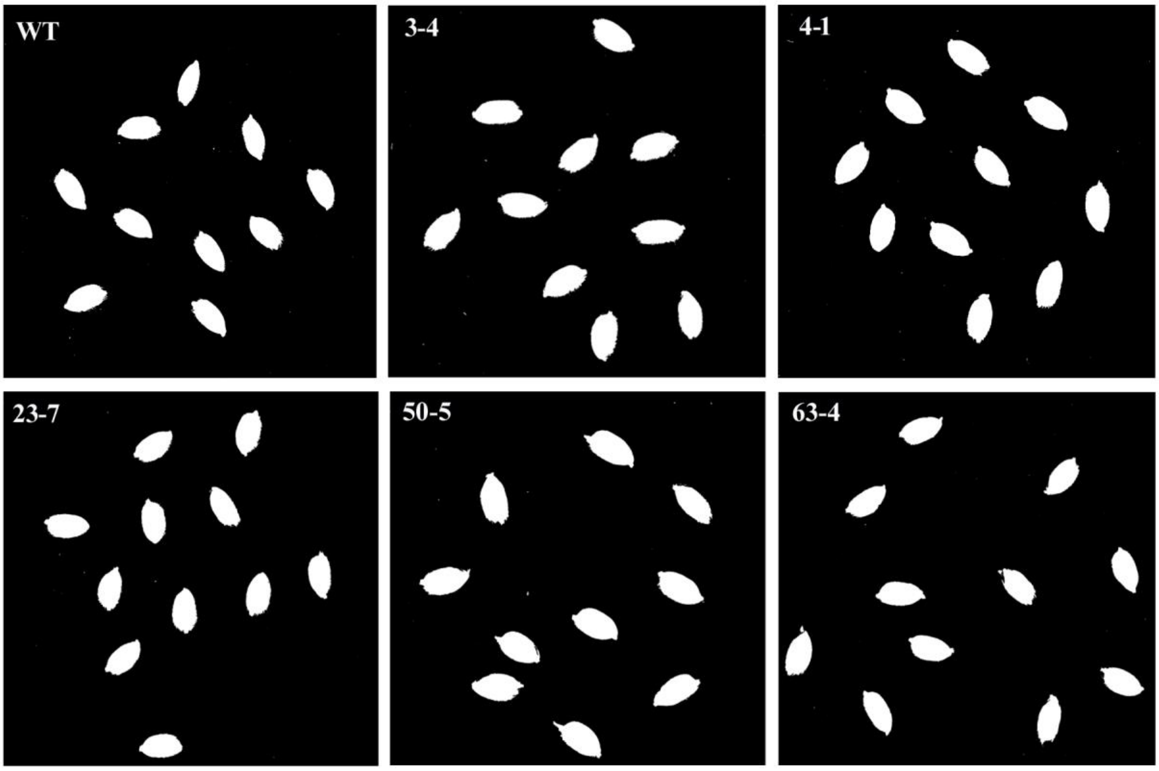

**Fig. S15: ImageJ-based analysis of seed size in wild-type (WT) and edited rice grains.** Representative binary images generated in ImageJ are shown, where individual seeds are isolated from the background for quantitative measurement of seed size. Identical image processing and analysis settings were applied to all samples to ensure accurate and unbiased comparison across genotypes.

**Table S1: Accession numbers of PEPC from different plant species**

| Sl. No. | Plant Species | Accession numbers |
|---|---|---|
| 1 | *Flaveria_trinervia* | UniProt Id- P30694 |
| 2 | *Sorghum bicolor* | Uniport Id- P15804 |
| 3 | *Zea mays* | Uniport Id- P04711 |
| 4 | *Panicum virgatum* | Phytozome genome ID: 450 • NCBI taxonomy ID: 38727 |
| 5 | *Panicum hallii* | Phytozome genome ID: 308 • NCBI taxonomy ID: 206008 |
| 6 | *Miscanthus sinensis* | Phytozome genome ID: 497 • NCBI taxonomy ID: 62337 |
| 7 | *Andropogon gerardi* | Phytozome genome ID: 783 • NCBI taxonomy ID: 79824 |
| 8 | *Gossypium hirsutum* | Phytozome genome ID: 458 • NCBI taxonomy ID: 3635 |
| 9 | *Hordeum vulgare* | Phytozome genome ID: 462 • NCBI taxonomy ID: 112509 |
| 10 | *Glycine max* | Phytozome genome ID: 275 • NCBI taxonomy ID: 3847 |
| 11 | *Solanum tuberosum* | Phytozome genome ID: 448 • NCBI taxonomy ID: 4113 |
| 12 | *Coffea arabica* | Phytozome genome ID: 453 • NCBI taxonomy ID: 13443 |
| 13 | *Oryza sativa* | Phytozome genome ID: 499 • NCBI taxonomy ID: 39947 |
| 14 | *Flaveria pringlei* | GenBank Id: CAA88829.1 |
| 15 | *Nicotiana tabacum* | Uniport Id- P27154 |
| 16 | *Arabidopsis thaliana* | Uniport Id- Q84VW9 |

**Table S2: List of primers used for this study**

| Sl No. | Primer name | Primer Sequence (5'-3') | Purpose |
|---|---|---|---|
| 1 | 574-OsPEPC_F | AGGTCGACCTCGAGATGGCGGGGAAGGTGGAGAAGATG | Amplification of PEPC from cDNA |
| 2 | 575-OsPEPC-R | CGGGGTACCTTAGCCGGTGTTCTGCATTCCAGCAGC | |
| 3 | 576-F1 | ATGGCATGCATCCACCAATTG | Site directed mutagenesis Arg>Gly (Invitro assay-PEPC kinetic efficiency) |
| 4 | 579-R2 | GGAGGCGAAGGCcCTGCTTCAGGTAGAG | |
| 5 | 580-F3 | CTACCTGAAGCAGgGCCTTCGCCTCCGC | |
| 6 | 706-PE2-F1 | AGGATCTAGCGGAGGATCCT | PCR for fragment generation from pK-PE2 and pH-ePPE2 |
| 7 | 707-PE2-R1 | TAGGTCTCCGCTGCTGCCGCCGCT | |
| 8 | 708-ePE2-F2 | CGGGTCTCACAGCGCCACAGTGGTGTCCG | |
| 9 | 709-ePE2-R2 | TCTCTAGACTCTCACACCTTCCTT | |
| 10 | 898-PEPC-DSF | ACACTCTTTCCCTACACGACGCTCTTCCGATCTGAAACTGAGGGCCAACTGTG | Deep sequencing |
| 11 | 899-PEPC-DSR | GACTGGAGTTCAGACGTGTGCTCTTCCGATCTCGGCTTAGACCAGTCCATGA | |
| 12 | 1150-PEPC-F | GAAACTGAGGGCCAACTGTG | Screening of mutants for PEPC target |
| 13 | 1151-PEPC-R | CGGCTTAGACCAGTCCATGA | |
| 14 | 1357-PEPC SEQ F | CCTTCAGGTGCGTCGACT | |
| 15 | 100-ZmUbi-Pr-F | ATGCTCACCCTGTTGTTTGG | Transgene free screening |
| 16 | 900-Cas9R_pGTR-en | AGCCTGGCAGAGAGAATAGC | |
| 17 | 788-HPT-F | GCTTCTGCGGGCGATTTGTGT | |
| 18 | 789-HPT-R | GGTCGCGGAGGCTATGGATGC | |
| 19 | 1386-RT_F1 | GCCACACTGCTGCCACTC | |
| 20 | 105-M13-R | CAGGAAACAGCTATGAC | |
| 21 | B58-ch12 F | CGTCCTCGTTGTCATCGC | Off-target analysis |
| 22 | B59-ch12 R | CGAAGGACCGTATCTGATCG | |
| 23 | B60-ch12 F1 | CATGGTCGTGGAGATCGCCGAG | |
| 24 | B61-ch12 R1 | TAGAACCATACACGTGGTCG | |
| 25 | B62-ch3 F | AGTGATAAAGGCGCGATGCA | |
| 26 | B63-ch3 R | GGACGGGGATCTTCCTGTGG | |
| 27 | B64-ch8 F | CCCATGGCTCTACTCCGAG | |
| 28 | B65-ch8 R | GAGGAATCCTCATCCGGGGA | |
| 29 | 834-OsActin-RT-F | CTCCCCCATGCTATCCTTCG | Expression analysis |
| 30 | 835-OsActin-RT-R | TGAATGAGTAACCACGCTCCG | |
| 31 | 836-Osbeta tubulin-RT-F | GGAGTCACATGCTGCCTAAGGTT | |
| 32 | 837-Osbeta tubulin-RT-R | TCACTGCCAGCTTACGGAGG | |

| | | |
|---|---|---|
| 33 | b12-RT-PEPC-F | TCTCAGCCACCAGACACAAT |
| 34 | b13-RT-PEPC-R | CCACAGCCATCTCATCAAGC |
| 35 | D56 NAS1 RT F | GTCTAACAGCCGGACGATCGAAAG |
| 36 | D57 NAS1 RT R | TTTCTCACTGTCATACACAGATGGC |
| 37 | D58 NAS1 RT F2 | GTTCCTGTACCCGATCGTC |
| 38 | D59 NAS1 RT R2 | CTTGTTGGCGGCAAACTCTT |
| 39 | D60 NAS2 RT F | TGAGTGCGTGCATAGTAATCCTGGC |
| 40 | D61 NAS2 RT R | CAGACGGTGACAAACACCTCTTGC |
| 41 | D62 NAS2 RT F2 | GTTCCTGTACCCGATCGTC |
| 42 | D63 NAS2 RT R2 | CCTCCATCTTGCAGCACTTG |
| 43 | D64 TOM1 RT F | AGTTGCAGATCGTATAGGGAGGAA |
| 44 | D65 TOM1 RT R | TCGGAAAATACATTTGGATATTGCT |
| 45 | D66 TOM1 RT F2 | TGATGAGGCCGGTGTTCTAC |
| 46 | D67 TOM1 RT R2 | ATGCACCAAGAAATCCAGCG |

**Table S3: Analysis of the potential off-target sites for *OsPEPC* sgRNA.**

| Potential off-target sites | | | | | |
|---|---|---|---|---|---|
| **Target** | **Sequence** | **Chromosome No.** | **Gene** | **Number of mismatches** | **Mutation detected** |
| Off-target 1 | CGTCGGCGTTGCGGAGGCGG **AGG** | 12 | OsKitaake12g123300 | 4 | None |
| Off-target 2 | TATAGCCATGGCGGAGGAGA **AGG** | 3 | OsKitaake03g365700 | 4 | None |
| Off-target 3 | CGGCGGCATAGCGGAGGCGG **AGG** | 8 | OsKitaake08g112366 | 5 | None |

**Sequence S1: pegRNA sequence with components**

pegRNA (R883G): Spacer-esgRNA-RT Template-PBS-Terminator

TGTAGGCATTGCGGAGGCGAgtttaagagctatgctggaaacagcatagcaagtttaaataaggctagtccgttatca
acttgaaaaagtggcaccgagtcggtgcacctgaagcagGgccttcgcctccgcaaTTTTTTT

**Sequence S2: epegRNA sequence with components**

epegRNA (R883G): Spacer-esgRNA-RT Template-PBS-Linker-tevopreQ1-Terminator

TGTAGGCATTGCGGAGGCGAgtttaagagctatgctggaaacagcatagcaagtttaaataaggctagtccgttatca
acttgaaaaagtggcaccgagtcggtgcacctgaagcagGgccttcgcctccgcaaAATAAAGGCGCGGTTCTAT
CTAGTTACGCGTTAAACCAACTAGAATTTTTTT
